# A Model for the Diversity Explosion Fundamental to Metastasis

**DOI:** 10.64898/2025.12.13.694153

**Authors:** Edward Tim O’Brien

## Abstract

Most deaths from cancer are caused by their metastases. Along with the ability to evade the immune system and to invade and disrupt very different tissue environments, metastases usually develop resistance to whatever therapeutic approaches are tried. Their extreme adaptability requires a diversity of traits from which natural selection can choose, but current models, such as the epithelial-to-mesenchymal transition (EMT), do not directly address how this diversity is generated. We observed that single cell clones of Panc1 human pancreas cancer cells that had had crispR knock outs (KO) of the gene for activin-like kinase 4 (ALK4) had developed a markedly diverse morphology. Time-lapse and fluorescence microscopy, FACS analysis and mitotic chromosome squashes provided evidence that the cells diversify profoundly in size, behavior and chromosome number. This diversity appears to develop through cytokinesis failure, the generation of very large multinuclear cells that exhibit a high range of nuclear configurations, coupled with continued cell divisions that sometimes generate three, four, and more daughter cells in one cell division. This astounding activity often resulted in cells that were much smaller than the normal Panc1 cells, and indirect evidence suggests they were significantly sub-ploidy, yet underwent regular cell divisions. A subset of the smaller cells were highly motile, and were observed to sometimes merge together or even into larger cells. This phenomenon could provide an unappreciated venue for transferring mutated genes, chromosome fragments, whole chromosomes or sets of chromosomes into an invaded cell. We suggest that these observations may be a serendipitous illustration of the missing link between the genetic mutations implicated in the development of primary carcinoma, and the extreme genomic diversity that characterizes malignant metastases.

**Graphical Summary:** Single cell clones of Panc 1 ALK4 KO cultures changed from largely uniformly epithelioid in character (A) to cultures composed of an unexpectedly wide variety of cells, and that this variety is not explained by the traditional “EMT” (epithelial-to-mesenchymal-transition). These cells range from very small to very large, have varying degrees of motility, chromosome number, and cell division behavior. We describe the new morphologies and behaviors, and present time lapse movies and microscopic evidence that help explain the evolution of this variety. These new cell types integrate into colonies where the smallest cells gather into spherical masses (D). We believe that the heterogeneity in genomic content and behavior provides the variety needed for natural selection to select cells that are resistant to most treatments for metastatic cancers.

## Introduction

For most cancers it is the metastases that lead to the death of the patient (Dillekås et al. 2019; Chaffer and Weinberg 2011). This remains true despite advances in chemotherapy, immune checkpoint inhibition, targeted therapies, and other innovations. Moreover, metastases often develop resistance to whatever regimen is initiated, in some cases exacerbated by the treatment itself, and often regrow after an initial positive effect (Chaffer and Weinberg 2011; Hata et al. 2016; Isozaki et al. 2023; Naxerova 2025b; Sequist et al. 2011). The ability of metastases to resist radio- and chemotherapy and to colonize new tissue environments indicates that the descendants of the founder primary tumor cells have diversified such that some variants survive treatment (de Bruin et al. 2014; Davoli et al. 2017; Francescangeli et al. 2023; Hata et al. 2016; Klein 2013; Shoshani, Bakker, et al. 2021; Trakala et al. 2021; Naxerova 2025a). Understanding how this diversification develops, and how diverse the descendants can be, is critical to creating new approaches to preventing metastasis in the first place and potentially for treating advanced disease.

The relationship of metastases to primary tumors has been actively investigated for centuries. In the later 1800s and early 1900s, Hansemann and Boveri each emphasized that the structure of nuclei became less organized and less like the parental tissue cells as cancer progressed to malignant metastases (Hardy and Zacharias 2005; Boveri 2008; Hansford and Huntsman 2014). Hansemann documented unequal cell divisions, and postulated that the unequal distribution of chromosomes led to cancer development, while Boveri demonstrated in sea urchin embryos that aberrant cell divisions could disrupt nuclear structure via mis-segregation of chromosomes to daughter cells. By altering the centrosome-to-nucleus ratio, he showed that any cell division into other than two daughters must result in altered chromosome numbers in the daughters (aneuploidy). While these observations and insights have largely been proved true, it was not clear in their discussions how mis-segregation of chromosomes was thought to fit in the progression from normal epithelial tissue to primary tumor and then to disseminated metastases.

In contrast to an emphasis on whole chromosomal changes, mutations or fusions of specific genes have been found to be strongly associated with specific cancers and even of stages within cancer progression. Beginning with the discovery of oncogenes and tumor suppressor genes in the 1980s, investigations into specific genes and gene networks have generated many new insights and therapeutic approaches. Activating mutations in genes and gene fusions that stimulate abnormal cell growth, such as HER2 in breast cancer, EGFR in lung, pax3/Fox01 in sarcomas, and H-, N- and K- RAS, and PIK3CA and others in gastrointestinal cancers, have been identified, and inhibitors shown to be useful in prolonging patient survival (Fan et al. 2021; Pandey et al. 2024). At the same time, inactivating mutations in genes that normally prevent unnecessary cell division, or prevent the division of cells with damaged or incompletely synthesized DNA, have also been identified. Inactivation or loss of ‘guardians of the genome,’ such as BRACA 2, TP53 and CDKN2A (p16) are found in most advanced cancers, but treatment approaches to restore guardian function have proven difficult to develop. Thus, in spite of the many successes and insights of the genetic approach, the dire consequences of metastases to the patient persist.

Changes in specific genes do correlate with an important event in metastasis: the epithelial- to-mesenchymal transition (EMT; **Figure 1A**). EMT studies focus on the loss of genes for E-cadherin, increases in the expression of proteases, and upregulation of transcription factors such as slug, snail, and twist (Boyer et al. 1989; Thiery 2003; Shibue and Weinberg 2017; Brabletz et al. 2018).

**Figure 1.**
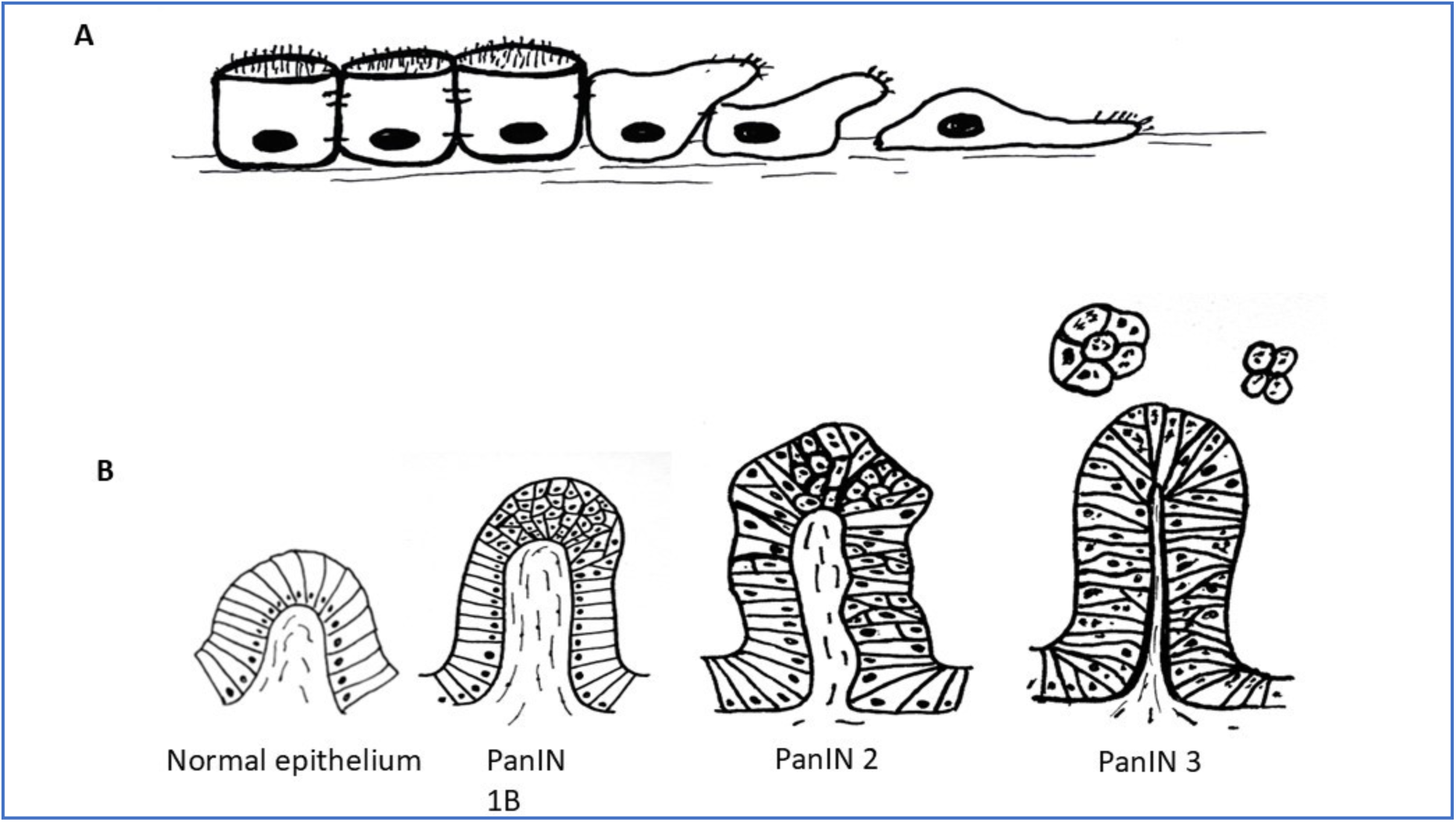
Drawings outlining the essentials of the epithelial to mesenchymal transition (“EMT”) (A) and a more representative depiction of the development of pancreas cancer from hyperplasia to metastasis (B). Drawing B is re-drawn for convenience from that shown in the Johns Hopkins University Pathology website: https://pathology.jhu.edu/pancreas/medical-professionals/duct-lesions, and is meant to Illustrate the enormous complexity that develops between an epithelium and metastatic cancer. (Please refer to the original source figure for a more precise illustration).

Migration, invasion, anchorage independent growth and ability to form tumors in mouse models are all increased as a cell transitions toward the mesenchymal state. Consistent with their importance in cancer, increases in measures of EMT has been correlated with changes in patient prognosis (Brabletz et al. 2018). On the other hand, the EMT model itself does not capture the hyperplasia driven by the driver gene mutations, or the nuclear and chromosomal diversity that is characteristic of advanced metastatic disease.

Pancreas cancer (pancreatic ductal adenocarcinoma, or PDAC ) is broadly typical of many solid tumors. As described above, a single cell in a well-organized epithelium undergoes a gain-of-function mutation in a Kras GTPase gene, which stimulates that cell to enter the cell cycle. The first stage in PDAC, where cells have undergone extra growth in a localized, polyp-like area, are still mononucleate and show normal mitotic figures, is termed a pancreatic intraepithelial neoplasia, or PanIN, stage 1A (Hezel et al., n.d.) (**Figure 1B-1**). In the progression to PDAC, statistical analysis of mutational timing strongly argue that the Kras mutation that drives the initial hyperplasia takes place on average about 11 years before diagnosis (Yachida et al. 2010; Iacobuzio-Donahue et al. 2012), arguing that that mutation by itself does not directly stimulate metastasis. The subsequent stages of PanINs show progressively greater tissue distortion and displacement, more cell divisions taking place away from the basement membrane, and a higher fraction of distorted and aberrant nuclei and mitotic figures as they progress (**Figure 1B -PanINs 2-3**). These changes correspond in time with sequential loss-of-function mutations in the guardian genes, p53 and p16. (Yachida et al. 2010; Iacobuzio-Donahue et al. 2012). Finally, mutations in the TGFbeta-targeted transcription factor SMAD4 is one of the last genetic changes in the progression to true PDAC. These latter loss-of-function mutations happen on average only a year or two before diagnosis, implying, at least in the case of PDAC, a more direct connection to metastatic behavior. The fact that PDAC is diagnosed most often when it is not resectable, and has a dismal prognosis due to early metastasis, argues that it might be useful to consider the later PanINs to be true PDAC, well along the pathway to metastasis.

Cell Biologists since the 1990s have argued that chromothrypsis, mitotic slippage, mitotic failure, cytokinesis failure and other mechanical events related to cell division would contribute to aneuploidy and chromosomal instability (CIN) and this could contribute to cancer development (Silkworth et al. 2009; Orth et al. 2011; Panopoulos et al. 2014; Shoshani, Brunner, et al. 2021; Herman et al. 2022; Krupina et al. 2024). At the same time, the pathway by which aneuploidy itself could promote and accelerate metastasis and generate the raw material that drives the evolution of resistance to treatment was still unclear. In this report we describe patterns of cell division and cell division failure that generate daughter cells with many different genomic and phenotypic properties. Some of these patterns have not to our knowledge been observed before in human cells. The fact that these patterns can take place at all, and the downstream consequences of these cell divisions, may provide the missing connections between the insights of Hannsemann and Boveri, known genetic changes, and the actual production of the diversity that allows metastases to evolve and adapt.

Our model is the Panc1 cell line, one of the models used by Zhang et al (2025) to test the effects of loss of the TGFbeta family member ALK4. Genetically, Panc 1 cells fall within that sequence of PDAC induction described above, in that KRAS is mutated, TP53 is partially inactivated, p16 homozygous deleted, and SMAD4 is present (Deer et al. 2010). Zhang et al had observed that loss of, or loss of heterozygosity of, the gene for the TGFbeta family member ALK4 was associated with poor prognosis in both pancreas and breast cancers (Zhang et al 2025). They wanted to know what behaviors and phenomena in vitro and in mouse models would correlate with the increased aggressiveness seen in patients. They generated crispR knockout cell lines of ALK4 in Panc 1 and other cell lines, and showed many of the patterns consistent with a metastatic state: greater EMT, increased anchorage independent growth, migration and invasion, and increased expression of EMT-associated genes. They went on to investigate the mechanisms driving these changes. At the same time, it was noticed that clones of the Panc1 ALK4 KO cells in particular exhibited unusual growth patterns in culture that pointed to a possible further diversification in cell morphology and behavior. Since ALK4 signals downstream through SMAD4, and SMAD 4 was the final change that seemed to correlate with full metastatic disease, we wondered whether the different morphologies we saw might be reflecting a later stage in the metastatic cascade, and whether this model could uncover clues to how the diversity in morphology had been generated.

## Results

### Knockout of ALK4 in Panc1 cells is associated with the appearance of several characteristic phenotypic and genotypic subtypes

Parental Panc 1 and empty vector (EV) control Panc 1 cell cultures showed a relatively characteristic epithelial morphology (**Figure 2; A,B**), retaining a generally epithelial morphology: monolayers with cuboidal or hexagonal packing with a low percentage of cells above the monolayer. In contrast, cultures of ALK 4 KO cells plated at the same time and cell density developed many apparently smaller cells on their apical surfaces, forming a second or even third layer. This pattern is readily apparent even at the low magnification, phase microscopic level used for daily cell culture (**Figure 2**). Parental (A;a) and Empty Vector (EV) (B,b) cells appeared largely epithelial in character (cuboidal or hexagonal shaped, closely packed and somewhat uniform) (**Figure 2A** and B). They had very occasional multinuclear (“MN”) cells and also had a few cells on the apical surface of attached cells, but these were not common. In contrast, the KO ALK4 cells showed a much greater diversity of form, yet the cultures grew in a particular pattern that was relatively consistent (**Figure 2 C-F**). Cells at the substrate surface were generally of medium size (≈20 µm diameter in the plane of the substrate). These medium-sized cells appeared by phase microscopy to be mononucleate, and the nucleus and cell body were similar in apparent size and morphology to the parental and EV cells. Sparsely (but consistently) distributed among the cells in the KO cultures were several types of multinuclear cells. An example of a large MN cell is shown in **Figure 2C** and c’. The diversity of MN cells is described in more detail below. Accompanying the EM and MN cells were cells that appeared smaller than either the MN or EM cells. These smaller cells appeared by phase microscopy to be of two general types: one that was spread and well attached to the substrate (smaller adherent, mobile, “SAM” cells), and one that appeared poorly attached to the substrate, were spherical, and gathered in mini-colonies on the apical surface of the monolayer. These spherical cells appeared to be less adherent to the substrate, but well attached to themselves and were somewhat adherent to the cells below them. We refer to these cells as “poorly adherent, small round” “PASR” cells. At the magnification of **Figure 2**, SAM and PASR cells are not easily distinguished, but the characteristic aggregations of apparently smaller, round PSAR cells are readily apparent, appearing as phase dense collections (C-F). These mini-colonies were very obvious and characteristic of the early passages of the KO cultures, and persisted into later ones as well. Individual cells within the dark masses of cells on the apical surface of the more epithelial cells are just distinguishable at this magnification, and are shown in more detail in later Figures.

**Figure 2.**
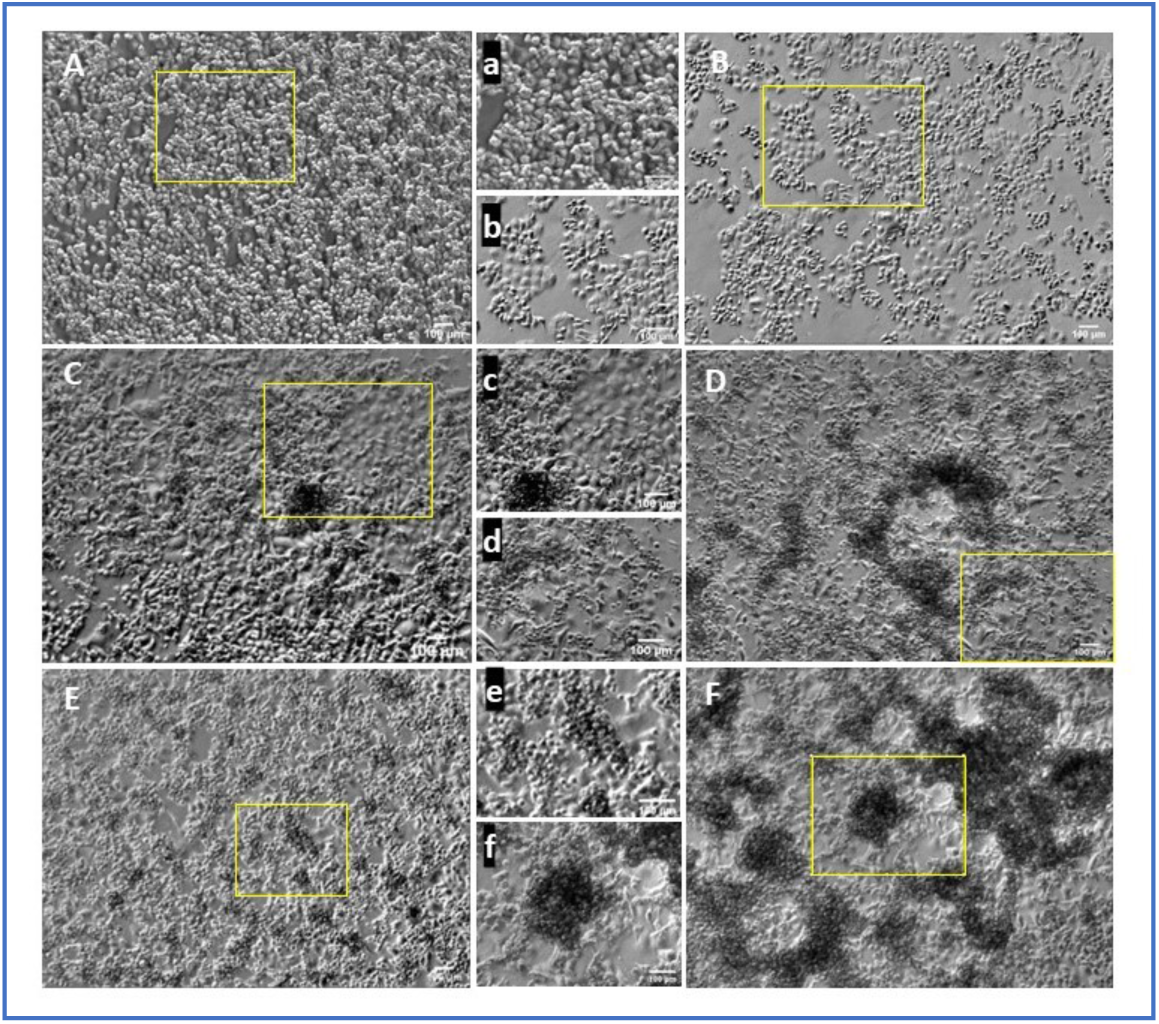
5X transmitted light views of Panc 1 parental (A;a), empty vector transfected (EV) (B;b), and crispR KO of ALK4 cell cultures (C-F) on plain tissue culture plastic. Routine cell culture imaging showed that parental (A) and EV (B) Panc 1 cells are somewhat diverse in morphology, but largely retained areas of epithelial morphology (illustrated in magnified areas (a,b)). They also exhibit a somewhat uniform range of sizes, with rounded, recently divided or mitotic cells on the apical surface of the epithelioid cells. In contrast, crispR KO Panc 1 cells (KO1 and KO2 (C,D) and KO3 (E,F)) commonly exhibited much greater morphologic and behavioral diversity: a greater frequency and variety of large multinuclear cells, and several types of cells that were smaller than typically seen in controls. One subtype of the smaller cells tended to accumulate in roughly spherical groups on the apical surface of the more epithelial, adherent cells. Examples of these accumulations are shown in E and F.

To see whether the morphologic differences we observed by phase microscopy were reflected in nuclear morphology, we evaluated images of DAPI-labeled nuclei from three experiments. These experiments were done to evaluate golgi morphology after ALK4 loss, and the DAPI channel was acquired for context. Without an F-actin or cytoplasmic membrane stain we could not confidently decide when several nuclei were in one cell, so each nucleus was evaluated on its own for size, and volumes were estimated as outlined in the Figure legend. **Figure 3** shows representative images from the 61 EV and KO cell images, where the diversity of cell morphology is reflected in a diversity of nuclear forms, including many smaller than normal nuclei in the KO cultures (3A). To compare the 3 cultures quantitatively, a histogram of nuclear volumes is shown in

**Figure 3.**
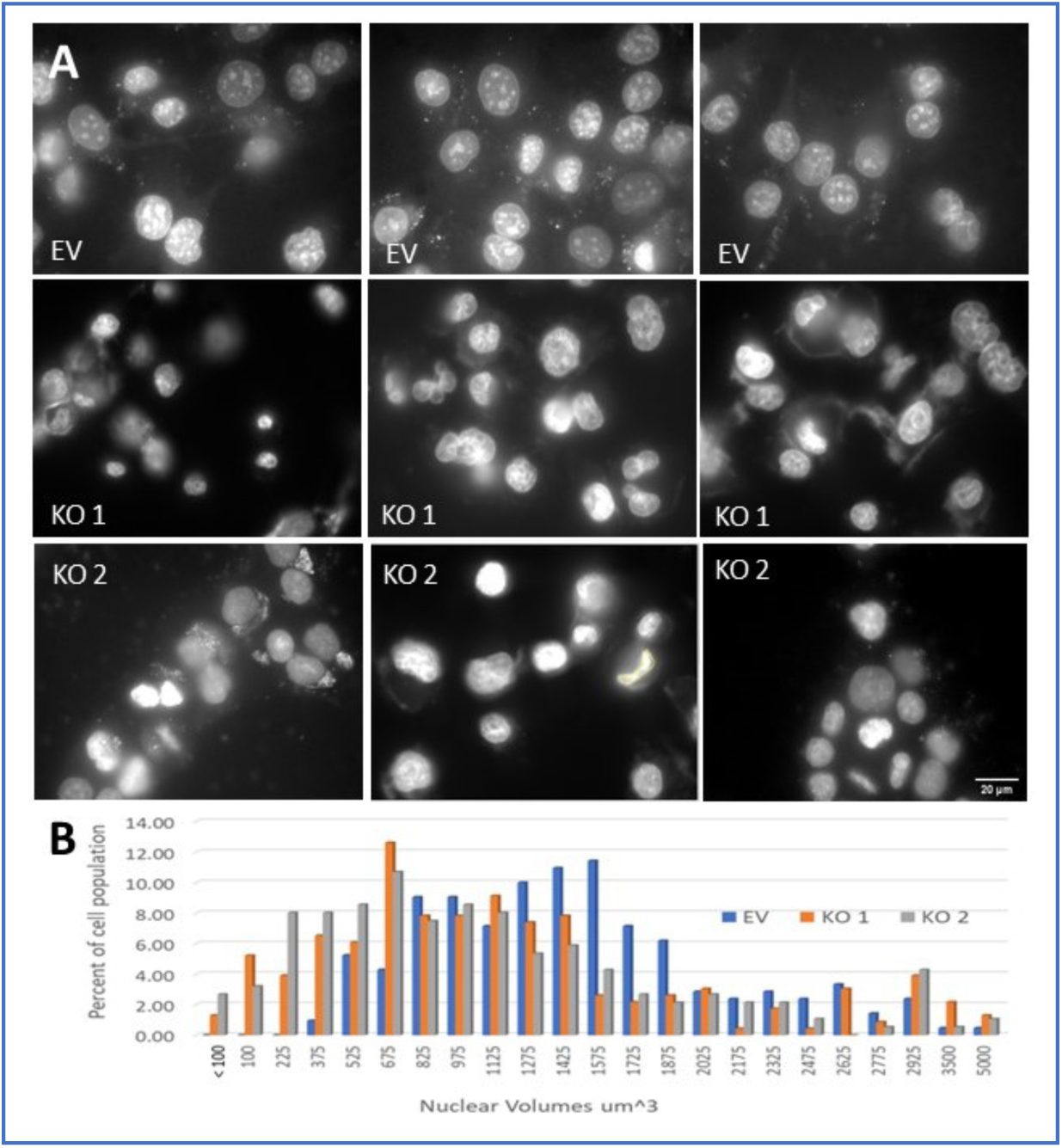
Nuclear volumes estimates of control vs crispR KO of ALK 4 cells show a much wider distribution of volumes in the KO cells. A.Representative widefield images from 3 experiments, showing DAPI fluorescence images of nuclei, and (B) tabulation of estimated nuclear volumes for ≈200 cells for each condition. Nuclear volumes were binned from 50 to 5000 cubic microns, for cultures transfected with empty vector (EV), and 2 independent crispR clones (KO). All images taken at 60X original magnification, and quantified as described in Methods.

Figure 3B, with a closeup of the histogram in 3C. Several focal planes in each field were obtained to assess the nuclei of cells attached to the substrate, and also to assess those on the next several layers of cells above the substrate. For a given cell type, nuclear volume is expected to be proportionate to DNA content. We therefore expect a randomly cycling population to show volumes that correspond to the 2N chromosome number (just after cell division) and a gradient up to 4N (tetraploid-just before cell division). The EV cells show a pattern consistent with this expectation, with volumes centering from 850 to 1600 μm^3^ and few cells below and above those values. In contrast, KO cultures showed a much higher proportion of nuclei much smaller than a normal 2N value, plus an apparent peak at about 3000 μm^3^.

To further characterize the high proportion of cells in ALK4 KO cultures with nuclei that were apparently smaller than normal controls, we used flow cytometry for DNA content with propidium iodide. This also showed a profound difference in the KO cell cultures. Cell cycle analysis using the FLoJo program (BD Biosciences), of empty vector (EV) and three ALK 4 KO cultures from 3 single cell clones, showed the presence of a prominent peak at about half the characteristic diploid (2N) chromosome complement of cells before S phase (G1) (Figure 4). Figure 4 shows two sets of comparisons between the EV (4A and C) and KO 1 (4B and D), and KO 2 and 3 (4E,F). Set 1 (A and B) and Set 2 (D-F) were done several weeks apart, and the settings on the FACS machine was slightly different. This resulted in the G1 peak being higher in the second set, but the pattern of observed differences remained. In both sets the EV cultures showed a classical cell cycle pattern of G1 (2N; diploid) to G2 (4N; tetraploid; just before mitosis) peaks with S phase (aneuploid) intermediate between them. Also observed was a shallow curve above G2/4N that represents DNA/cell greater than tetraploid. In contrast, the KO cultures (B, D-F) each showed a 2N peak at the same intensity value as the controls, but also showed an additional prominent peak at approximately one-half the G1 value seen in the controls (see red arrows in Figure 4 **B,D-F**). These peaks, at approximately the haploid (1N) value of DNA content/cell, suggest that a population of cells have evolved that have a significantly reduced number of chromosomes per cell. The position of this peak at about the haploid chromosome number could suggest that a population of cells has lost one set of parental chromosomes, and thus was actually haploid. Further experiments argue against this possibility. The fact that three, single cell ALK4 KO clones each showed the “haploid peak” suggests that this development may be an important characteristic of ALK4 loss, and is consistent with the distribution of nuclear volumes described in Figure 3.

**Figure 4.**
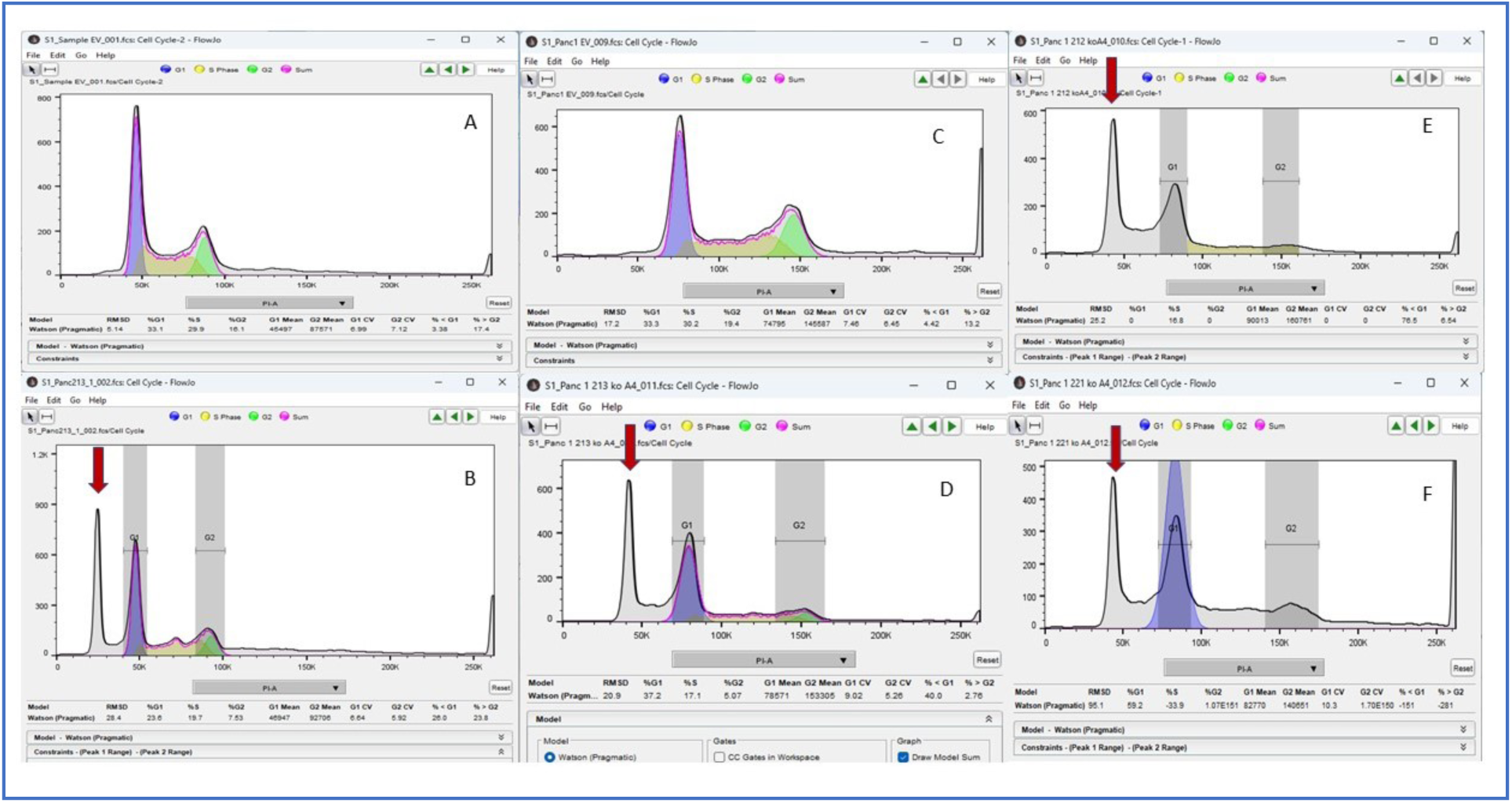
DNA content of 10,000 control (EV) and ALK 4 KO cultures by flow cytometry show anomalous peaks at ≈half diploid in KO cultures. Same day comparisons between EV and KO cultures are shown. For easy comparison A and B are shown in vertical pairs, as are C and D. The red arrows show the unexpected peaks in the KO nuclear DNA content distributions. Cultures were grown under normal conditions on tissue culture plastic to 60-80% confluence, washed in PBS and fixed with 70% ethanol, and prepared for DNA/cell cycle analysis with propidium iodide as described in Methods.

While the presence of the haploid peak seems to be the most dramatic difference between control and KO cells, the analysis itself cannot distinguish whether these sub-diploid cells are themselves cycling. In Figure 4 we emphasized the similarity in the G1 peak position in EV to that of the KO cultures by constraining the G1 to the same position in the KO samples as the EVs. This makes sense, since each set of EV and KO Panc 1 cultures were processed at the same time, with the same reagents. The gray sub-diploid peak thus stands out as very anomalous. Analyses done without such a constraint, both in FloJo and in ModFit (Verity Software House; Cytonome Verity, LLC) are presented for completeness for the first comparison set in **Supplemental Figure 1**. Both software algorithms model the KO cell populations as though the earlier, haploid peak is G1, and all the cells between that peak and the G2 are in various stages of S phase, and therefore cycling. This of course does not prove the sub-diploid cells are cycling normally, albeit with fewer chromosomes, but it is consistent with that interpretation and with the observations that follow (below).

To pinpoint the key changes underlying the sub-ploidy population and nuclear differences seen in the ALK4 KO cultures, we carried out chromosome squashes. We tried several protocols to observe banding using Wright and Geimsa staining, with limited success. Fortunately, the mounting medium in all cases contained DAPI, and we were able to obtain reasonable quantification of chromosome number in most experiments. The exact number of chromosome pairs in the squashes became more difficult to determine with increased chromosome number (≈70-100+), but, for the argument we are making, slight differences at high numbers are less consequential.

The squashes with low numbers were much more easily quantified. Of the 100-200 images taken, only 81 were sufficiently spread and good enough quality to be counted. The main criterion was that they were isolated enough from other squashes that they were not likely to have been scattered from a nearby squash. Representative images of counted squashes are shown in Figure 5, along with histograms of the number distributions of the chromosomes/squash. The histograms are displayed with bin widths of 5 (5B), or 1 (5C). As shown in the distributions, there was a preponderance of squashes with a very low number of chromosomes. Bin 1 in Figure 5B showed that almost half of the quantified squashes were present with 1-5 chromosomes. Figure 5C shows a more exact representation, with squashes having 1 or 3 chromosomes being most prevalent.

**Figure 5.**
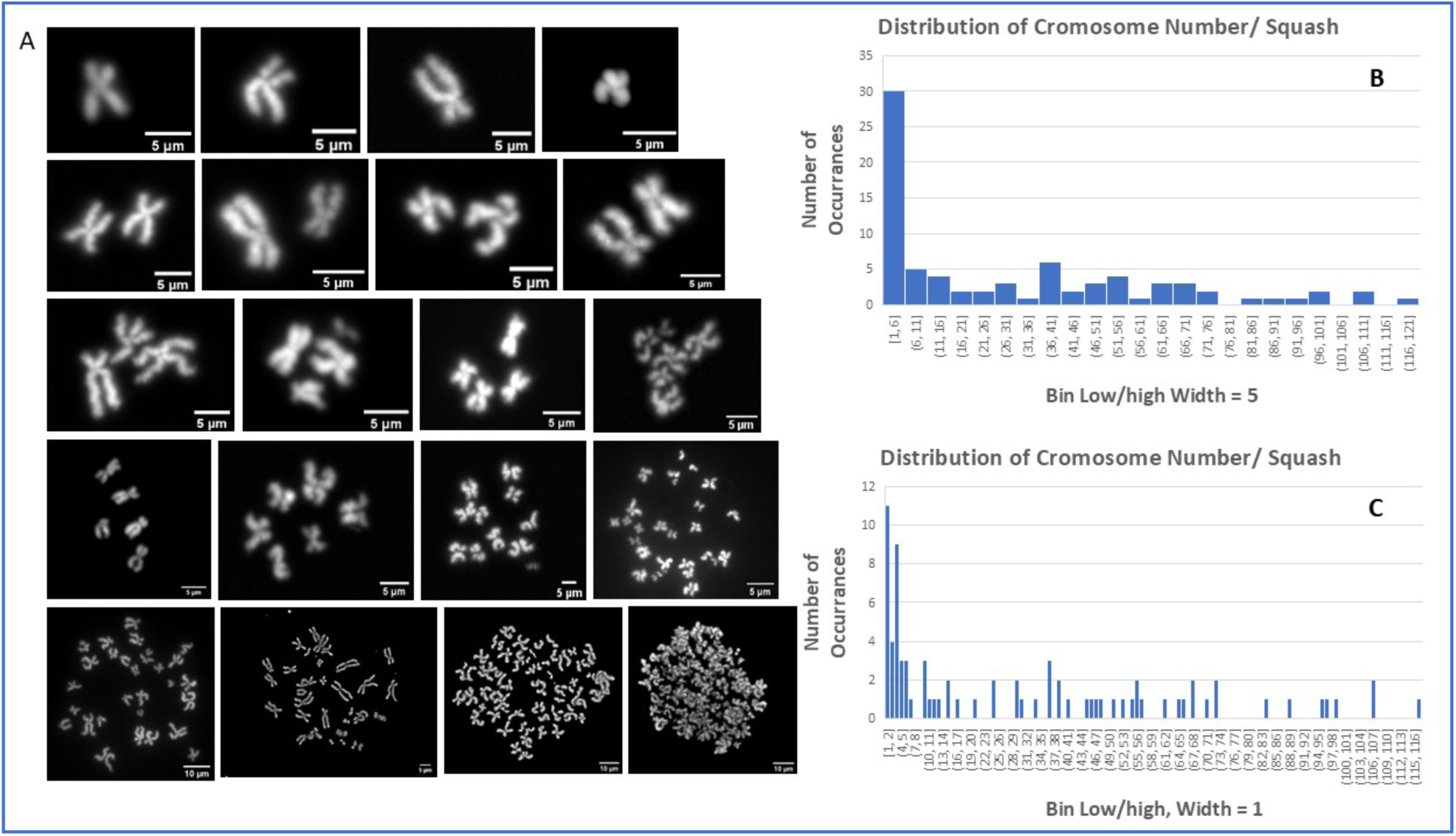
Chromosome number per cell of ALK 4 KO cells. Chromosome squashes of ALK4 KO cells showing examples of fluorescently labeled (DAPI) chromosomes prepared after blockage in metaphase. The histograms show the number of times a particular number of chromosome pairs were found (n-77). The same data is binned as 5 (B) or 1 (C) chromosome/cell.

It is interesting to compare these results with the cytometry data in Figure 4. That experiment showed an extra sub-ploidy peak at about ½-normal, or about 23 chromosomes. We assumed at first that this implied that these cells would be haploid – perhaps losing one parental genomic complement. In contrast, the karyotype results strongly suggest that this cell population is comprised of cells with very few chromosomes. One way to reconcile these findings is to observe that a high fraction of the cells with few chromosomes seem to have those with the longest, lowest number (Figure 5A). Chromosomes are numbered by mass or length, with longest being 1. Human chromosomes 1-7 combined contain ≈50% of the mass of the normal 46 pr human karyotype (Piovasan 2019), while 8-23 combined constitute the other half (with slight adjustment made for and X or Y chromosomes depending on the sex of the individual). A cell with a chromosome complement of just numbers 1-3 would comprise about 25% of normal diploid DNA mass. Thus the cells with fewest but longest chromosomes could provide a disproportionate fraction for the haploid peak. Further work will be necessary to determine whether such cells may represent an asymptote toward a “minimum genome necessary for survival” chromosome number, or at least minimum genetic complement. At present, the crucial and most interesting finding is that cells with low and very low numbers of chromosomes seem to exist, while culture growth and time-lapse movies argue that they are very active and able to reproduce (see below). These cells may represent a closer approximation to a diverse metastatic phenotype than simple EMT, and may have been relatively understudied thus far.

### Cell behavior of the ALK4 KO cultures

We wondered whether the categories of cellular size, shape, and genome content we observed would correspond to characteristic differences in behavior, and if the observed behaviors would offer clues to how this diversity, especially of the small, apparently sub-diploid cells, was being generated. To minimize the effects of passaging techniques that preferentially lose poorly adherent cells, and to promote reasonable spacing between cells, we used 8 µm pore transwell filters (PC membrane; Corning Life Sciences). Zhang et al. established that the ALK4 KO cells were faster at traversing these filters, either with or without a Matrigel (Corning Life Sciences) barrier (Zhang et al 2025). By using inserts that were one size smaller than our 12 well plates, we could place the filters directly onto the tissue culture plastic. In this way cells that traversed the pore could readily attach to culture plate. To increase the likelihood of attachment, the plates were briefly coated with a 10 g/ml solution of Matrigel (Corning Life Sciences; no growth factors) in PBS for 15 min, then the solution suctioned off and allowed to dry. Cultures were trypsinized and ≈20K cells were placed on the filter, normal medium (10% serum) placed on both sides of the filter, and incubated for 48 hrs. The filter was then removed, and cells allowed to grow for 10-14 days before placed in the time-lapse microscope system. Figure 6 shows a still from the movie presented in **Supplemental Movie 1**, with labels of the general classes of cells that appear to have differentiated from the single cell PANC1 ALK4 KO cell clone. The overall impression of these colonies is that the diversity observed in Figure 2 is preserved: an attached epithelioid cell layer is overgrown in places by masses of smaller cells. The roundish, large aggregates appear dark in the movie due to being many cell layers deep and above the primary plane of focus. The cells comprising the aggregates appear in the movie to be actively moving, and many are dividing. The tendency of these cells to collect into ≈3-dimensional forms suggest that the component cells have an appreciable affinity for sister/daughter cells, and possibly less for the planar substrate. Surrounding the aggregations appear to be several additional cell subtypes. There are still representatives of the “normal” Panc1 epithelioid cells present at the surface. We label these “EM” cells for epithelial-(presumably) mononuclear. A few larger cells are also present, labeled “MN” for (presumably) multi-nuclear. (At this magnification, multinuclearity is not clear, but further imaging will establish the likelihood.) The remainder of cells that can be seen moving rapidly between aggregates we term “SAM” cells, for <u>s</u>mall, <u>a</u>dherent and <u>m</u>otile. As observed in S**upplemental Movie 1**, these cells move rapidly, and change direction often. An exception are those cells that appear to follow along a track between aggregates. Here two lines of cells locomote in opposite directions, surprisingly reminiscent of worker ants. These cells tended to move along inter-colony paths for many hours, and occasionally would round up and divide, then rapidly resume locomotion. The cells along the linear paths could be tracked for an average of 6.2 hours (SD = 3.26), wherein they maintained a translocation rate of 19.3 µm/hr; SD = 6.99; n = 11). This rate is close to that observed for individual fibroblasts in vitro (Park et al. 2005; Wu et al. 2012) and also to the rates predicted by modeling fibroblast motility (Dokukina and Gracheva 2010), of between 6 and 30 µm/hr.

**Figure 6.**
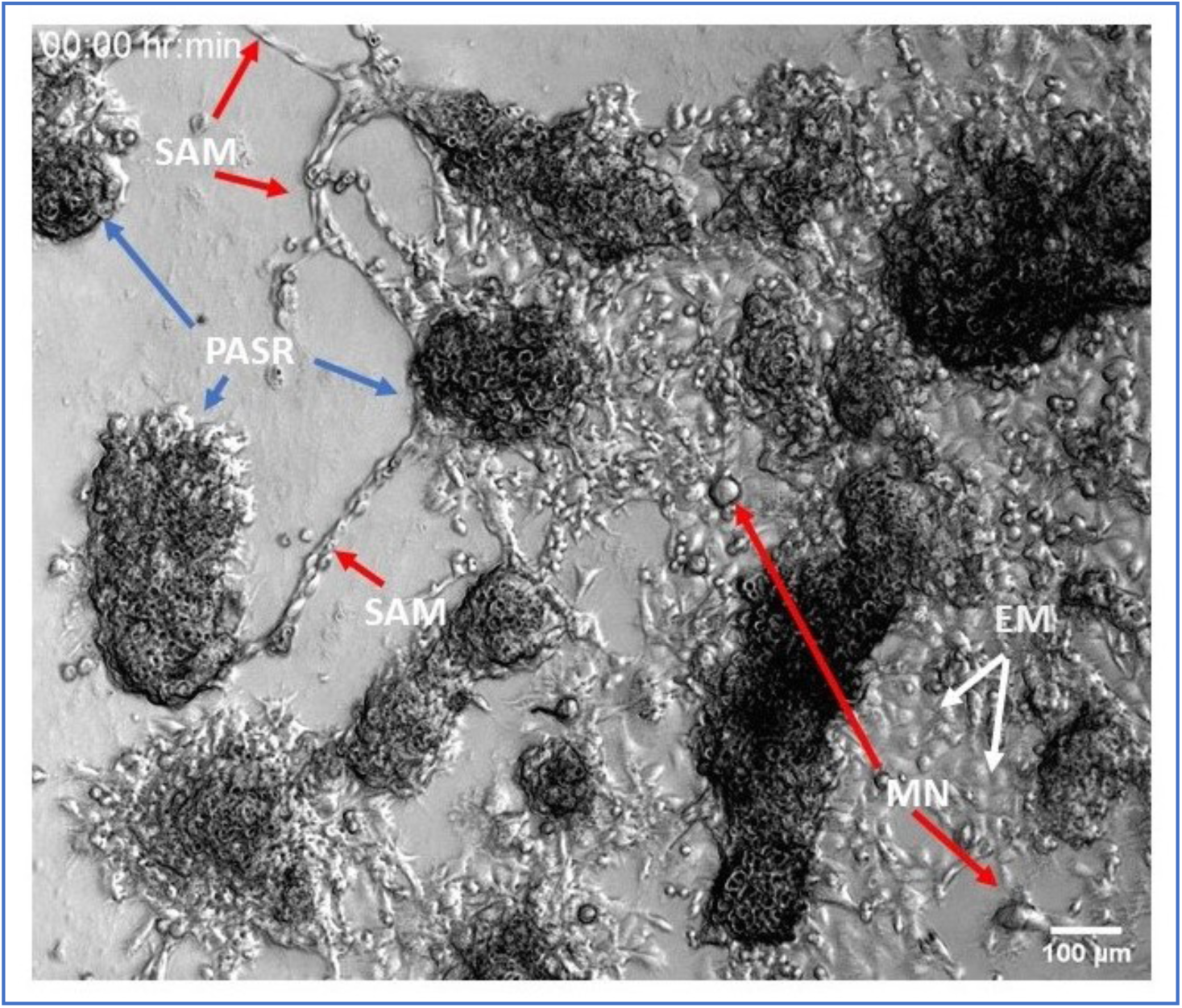
Still image of from Supplemental Movie 1. Panc 1 ALK4 Ko cells grown from a single cell clone, show dramatic diversity in morphology and behavior. Cells were plated on tissue culture plastic in a 12 well plate. The plate was treated with a scant coating of Matrigel, on an 8 µm transwell filter that had been placed directly on the plastic surface. The filter was removed after 24 hrs, and cultures allowed to grow undisturbed except for refeeding for 7-14 days. Images were acquired every 4 min in 5x phase mode with the Zeiss Live Cell station, with CO2 and 37 deg warming.

Interestingly the rates we observe were closer to that shown by Park et al for 3T3 cells that had been stimulated by HRAS. Thus, whether smaller physically, or containing fewer chromosomes than normal, the SAM cells appear to locomote at rates similar to full sized, mouse or human fibroblasts.

We wondered whether the cells that appeared smaller than the EM cells could be divided into separate subtypes. To determine whether these cells were actually distinguishable as subgroups on the basis of morphologic and behavioral characteristics, we measured the sizes of these cells when plated normally, without filters. We compared Panc 1 EV, ALK4 KO1 and cells from a parallel ALK4 KO culture plate that had been gently “rocked” for a minute (without spilling the medium), the media collected, the cells centrifuged down, and then the cells plated in parallel with the other cultures. In this way we could ascertain whether the poorly adherent cells were viable, and would develop much differently than the standard ALK4 KO cultures. To provide a standard metric of cell volume, we played the videos of the cells frame by frame, backwards and forwards, and determined the diameters of each cell that underwent cell division, just before cytokinesis started. This is the most spherical and greatest volume that cell would occupy. To minimize confusion, MN cells that divided into more than 2 daughters were not included and are considered later. As shown in Figure 7, we observed a large differences in the cell volume of populations between EV and both the KO parental and the ‘rocked off’ KO cultures. To our surprise there was no obvious subgrouping of the smaller cells in the KO cultures. Instead, at the scale we were able to measure, they appeared to be present in a gradient between smallest and cells of more EV size. Consistent with this result, the poorly adherent cells from KO cultures, rocked off gently from a parent KO culture, produced a distribution of cell sizes very similar to that of the parent culture (compare Figure 7 **B and C**). The Panc 1 EV cells were not perfectly uniform (Figure 7A and D), but the pattern of size distribution was markedly different from the KO cultures. Primarily, the EV cells were more than two-fold larger in volume, on average, than the KO cells. The average volume of the EV cells was 17,087 μm^3^(SD = 8259), and almost no cells were observed below 8000 μm^3^. In contrast the ALK4 KO cells had a preponderance of cells below 8000 μm^3^, and an average volume of 7657 μm^3^(SD = 7123). (For reference, the diameter of a cell with 8000 μm^3^ is 4.8 μm^3^, while that of 4000 μm^3^ is less than 4 µm.) Yet these cells were able to regrow a culture with a very similar population distribution to the parent. Interestingly, amid the ≈280 cells analyzed, almost no cell divisions resulted in more than two daughters. This suggests that, once formed, these smaller cells may maintain a stable genotype as they reproduce. This is in contrast to cell divisions observed in MN cells, which often resulted in more than two daughters (described below).

**Figure 7.**
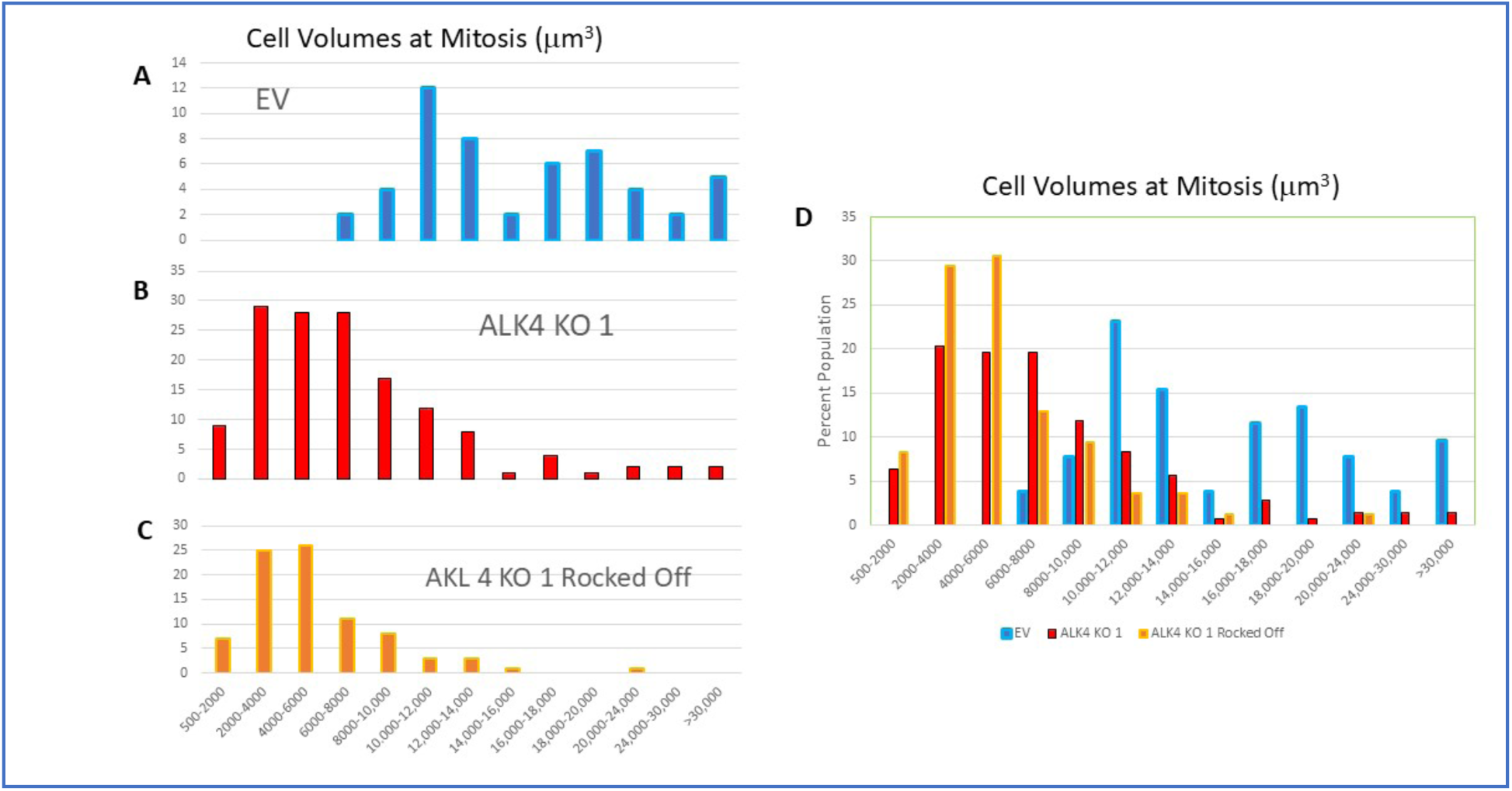
Comparison of cellular volumes between Control and ALK 4 KO cells at mitosis. When entering mitosis cells have maximum volumes, have rounded up, slightly detached from the substrate and therefore can be modeled as spherical. Cells cultures of equal confluence (60-80%) were imaged at 4 min intervals for 24 hr. Videos of cell cultures were observed both in forward and reverse to determine the last frame before cytokinesis. The diameter of each was then measured using ImageJ and volumes calculated as 4/3p(d/2)^3^. Consistent with the distributions of nuclear sizes, we find that the KO cell population were skewed toward the very small. Both normally passaged KO cultures and those regrown from cells gently rocked off from that culture showed very similar distributions. N=50 (EV), 144 (ALK4 KO), 86 (Rocked Off). A-C: Data plotted for individual cultures, D: Same data plotted together.

Although the smaller cells did not divide neatly into easily defined subgroups by maximum volume, they did follow some general trends that seemed consistent. These traits have obvious implications for the dissemination of cancerous cells during metastasis. First, the smaller cells appeared to be less adherent to the substrate than the larger EM and MN cells, especially during and just after cell division. As mentioned above, adherent cells at their maximum size and with all DNA replicated, round up and de-adhere slightly from the substrate surface before dividing. In a healthy epithelial tissue, under the signaling controls typical of that tissue, both daughters would then spread into the space vacated by the parent. Many cultured cancer cells also obey this generalization; even our Panc 1 EV cells would usually re-spread successfully and thus remain as a monolayer. This is evident in the lack of spherical cells building up in the EV cultures (Figure 1). In the ALK4 KO cultures, the smaller cells often de-adhered from the substrate, usually after rounding up for, or just after, cytokinesis. An example of very small cells exhibiting this behavior is presented in **Supplemental Movie 2**. These ALK 4 KO cells were separated from the main culture by allowing them to pass through a 3 µm transwell-type filter onto a scant coating of matigel on tissue culture plastic. They exhibit good motility until they round up for cell division. Just after cytokinesis, they fall off from the substrate and drift away. This is very typical.

Interestingly, we often observed de-adhered cells floating down into the field of view, and usually onto other, spread cells. The cells often piggyback on the larger cell, often for many hours. An example of this behavior is found in many of the movies presented, but in particular. **Supplemental Movie 3.** This also illustrates the directional motility of a subset of the multinuclear cells. This locomotion, probably of a binuclear cell, is very reminiscent of the keratocyte motion modeled extensively by Theirot (Labuz et al. 2023), and may be of interest to the biophysical community. Two small cells land on top of the large cell at about hour 4, and stay on top for 9 hours before one cell gains traction on the substrate and moves away. It then moves back onto the large cell, and the second cell moves off, before finally both small cells move away. This illustrates that the small spherical cells being carried are capable of individual motility. In terms of metastatic distribution, both the de-adhesion from an ECM while effectively binding to and to ride along with larger, more aggressive cells, would contribute to longer range movement of these putative cancerous seeds during metastasis.

The small cells often demonstrated a more troubling behavior: plunging in, under, and through other, usually larger cells. This fascinating behavior could be analogous to invasion into new tissues during metastasis and shows that a proportion of the very smallest cells have retained cell-cell recognition, adhesion and motility properties necessary for invading other cells, and the genes and regulatory circuitry to use them. An example of this behavior is shown in **Supplemental Movie 4**, and also in several later movies. Merging between recent daughter cells is also often observed, but it is difficult to distinguish that phenomenon from a failed separation after cytokinesis.

Genomic degradation and a progression toward the smallest cells from cells with normal ploidy appear to evolve from progressively aberrant multinuclear forms. Examples of the many configurations of nuclear forms we observe in multinuclear ALK 4 KO cells are shown in Figures 8 and 9. Figure 8 shows examples of the same scene from both phase contrast and fluorescence imaging modalities. We wanted to follow the changes in nuclear forms in time lapse, using Hoesch stained DNA (blue color in Figure). We found that the cells only tolerated 5-10 exposures at minimal exposure time, before senescing. We tried many DNA stains, of many wavelengths including far red excitation, and found the cells would still rapidly stop cycling and stop moving.

**Figure 8.**
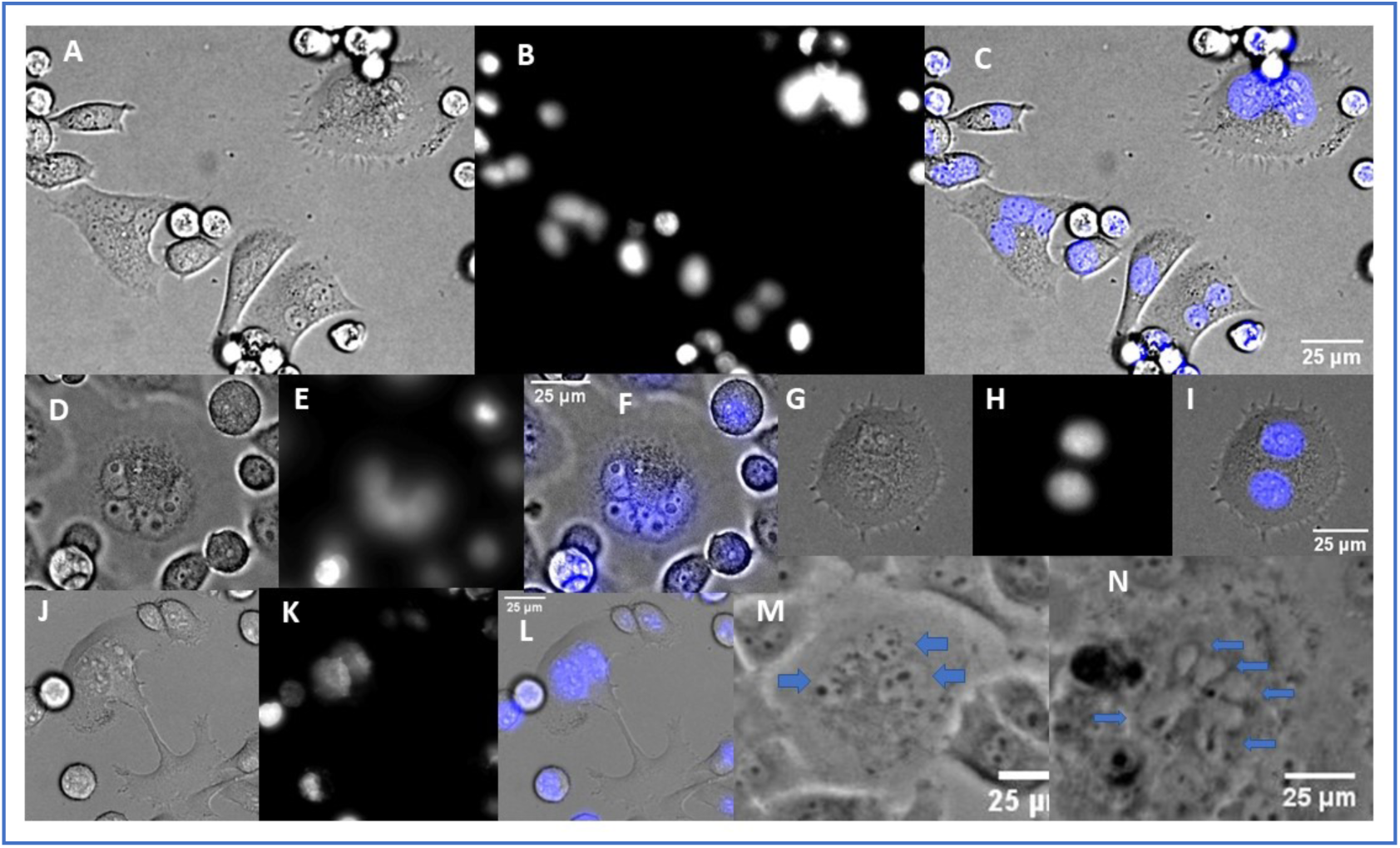
Live cell images of multinuclear cells using both fluorescence and phase light microscopy. The first frame of a movie of Hoesch stained cells planned to follow DNA through several cell cycles, is shown with and without the phase image. Later frames showed movement in these cells completely blocked. Images chosen to show examples of the variety of nuclear formations commonly encountered and their appearance in both modalities. Panels M and N show only the phase image, with blue arrows to indicate the very small nuclei present.

This strongly suggests that not all checkpoints have been abrogated in these cells, and they are very affected by reactive oxygen species or perhaps certain wavelengths of light. The images in Figure 8 **(A-L)** were taken from the first frame of the Hoesch stained movies, and illustrate how the fluorescent nuclei correlate with the phase images. This will be useful in interpreting several later movies without fluorescence. Images Figure 8**, M and N** shown without the nuclear stain, illustrate cells in phase contrast that have many smaller nuclei, like those seen in Figure 8 D-F. (See blue arrows in the unstained cells, (**Figure M and N**). Several similar nuclear configurations are shown in fixed, widefield images in Figure 9. F-actin, DAPI only, and the merged color images shown for context. Other configurations, including one very large nucleus, are also observed, but the examples of 2,3,4 and “many” are a sample of the variety commonly observed.

**Figure 9.**
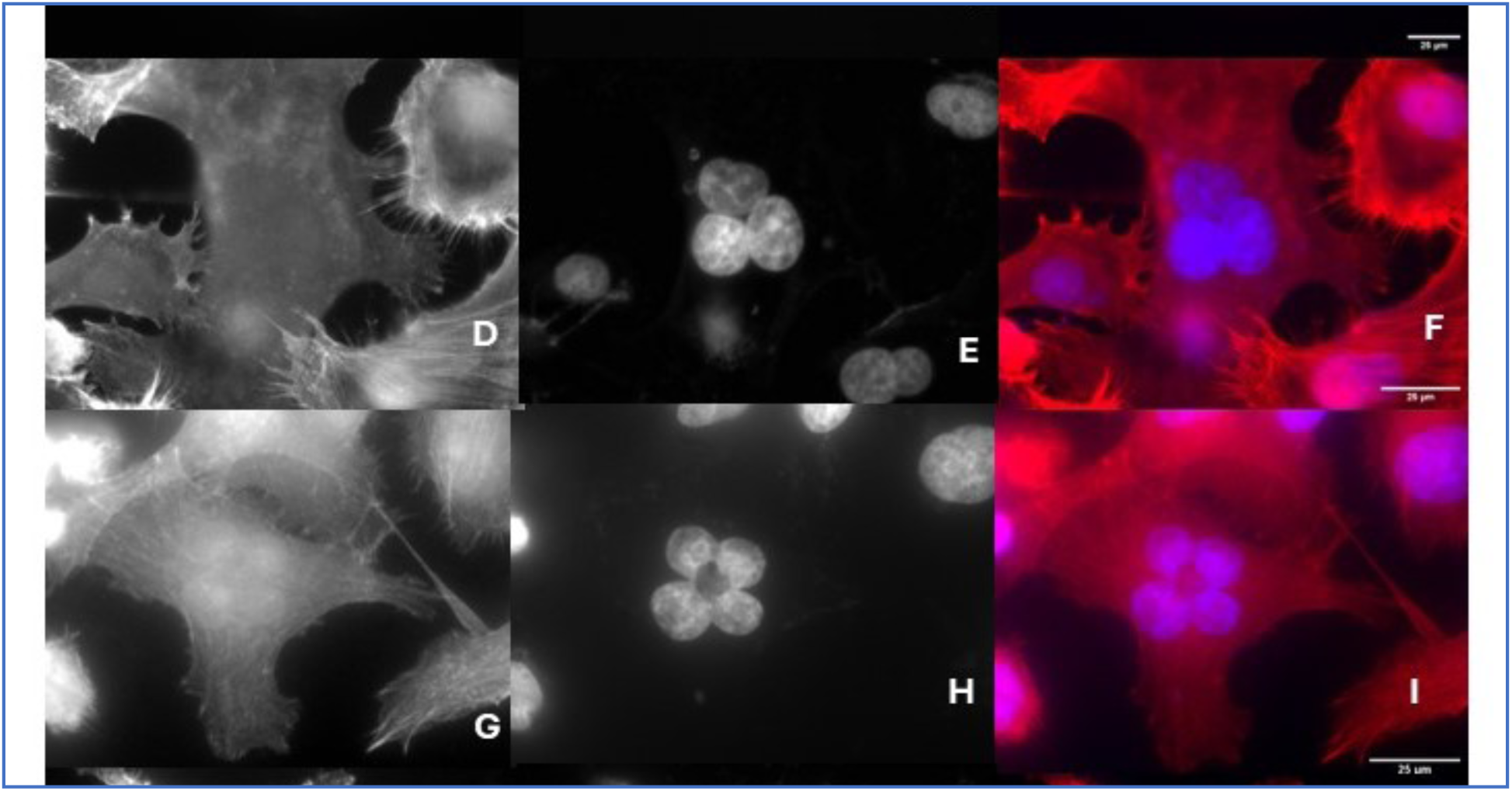
Fixed cell examples of nuclear conformations in ALK 4 KO multinuclear cells. Stains for actin (phalloidin; Figure 9A,D,G, and J) and DNA (DAPI; B,E,H,and K) are combined in the third column to show the nuclei in context. Several other conformations, such as very large, single nuclei are seen, but are less common.

Time lapse observations of ALK 4 KO cultures provide clues to the formation of extremely aberrant nuclei in the multinuclear cells. Disruption of the genome from the normal 46 pairs/cell appears to take place in this model system via (1) failed cytokinesis, (2) cell division greater than 1:2, (3) merging between cells, and (4) a very unusual breaking up of very large multinuclear cells into many small cells. This latter phenomenon we describe as “hatching.” **Supplemental movie 5** shows an example of failed cytokinesis, where a cell rounds up, stalls for several hours, and then respreads and moves away without dividing, thus becoming bi-nuclear. Importantly, after rounding up, the cell took ≈5 hrs to organize its chromosomes into a metaphase configuration. In metaphase, it spent another ≈3 hrs to begin anaphase, and then only 2 hr before it began to re-spread and move as one cell. Figure 10 shows frames from the movie, to illustrate these time points. For comparison, the time from rounding to cytokinesis in smaller ALK 4 Ko cells is about 1 hr. The extraordinary time spent in each phase indicates that the cell cycle apparatus was active, along with checkpoints that would normally assure proper chromosome segregation. But signals to prompt these cells to undergo apoptosis or senescence are not effective, and the cell survives and resumes normal motile and even further cel division events.

**Figure 10.**
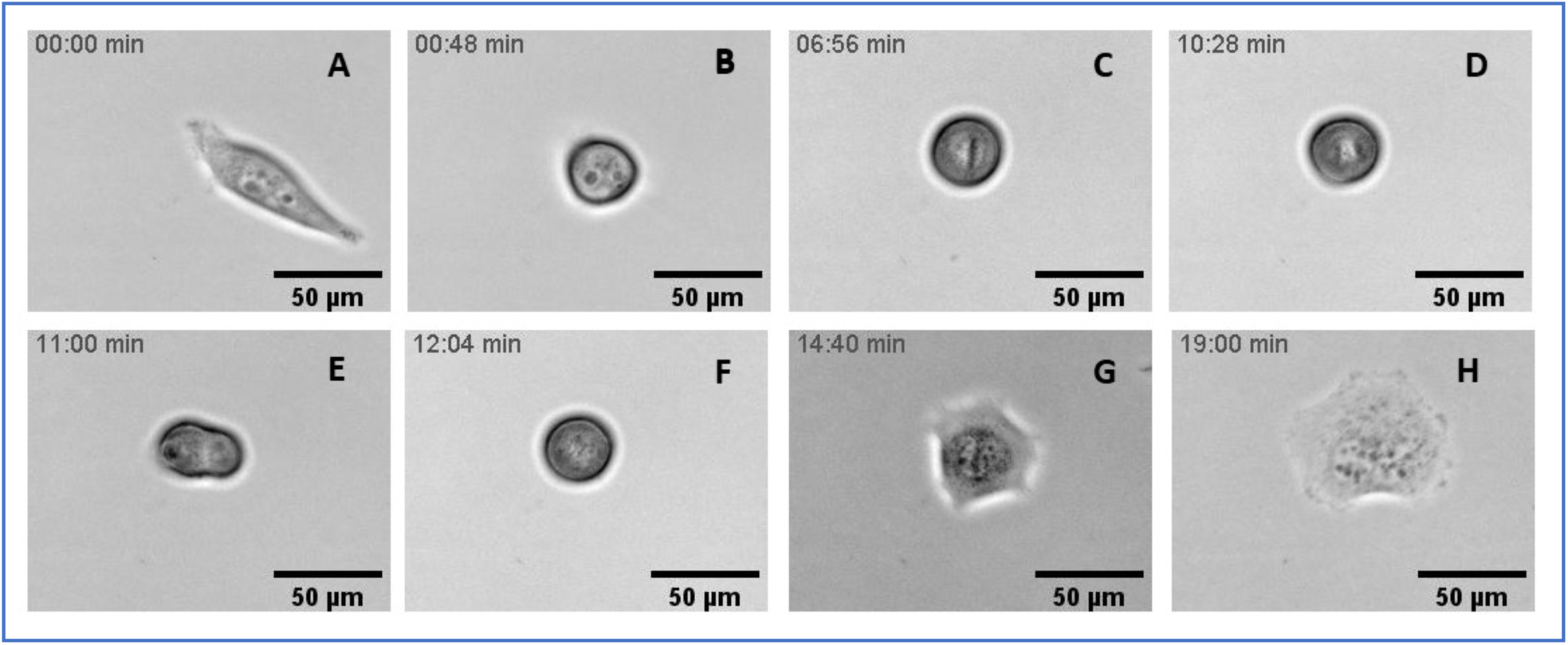
Frames from Supplemental Movie 5: an ALK 4 KO cell undergoing failed cytokinesis. Relative times in the sequence are seen in the upper left of each frame, illustrating the long periods that take place between cell cycle stages before the cell moves along as one binuclear cell.

**Figure 11.**
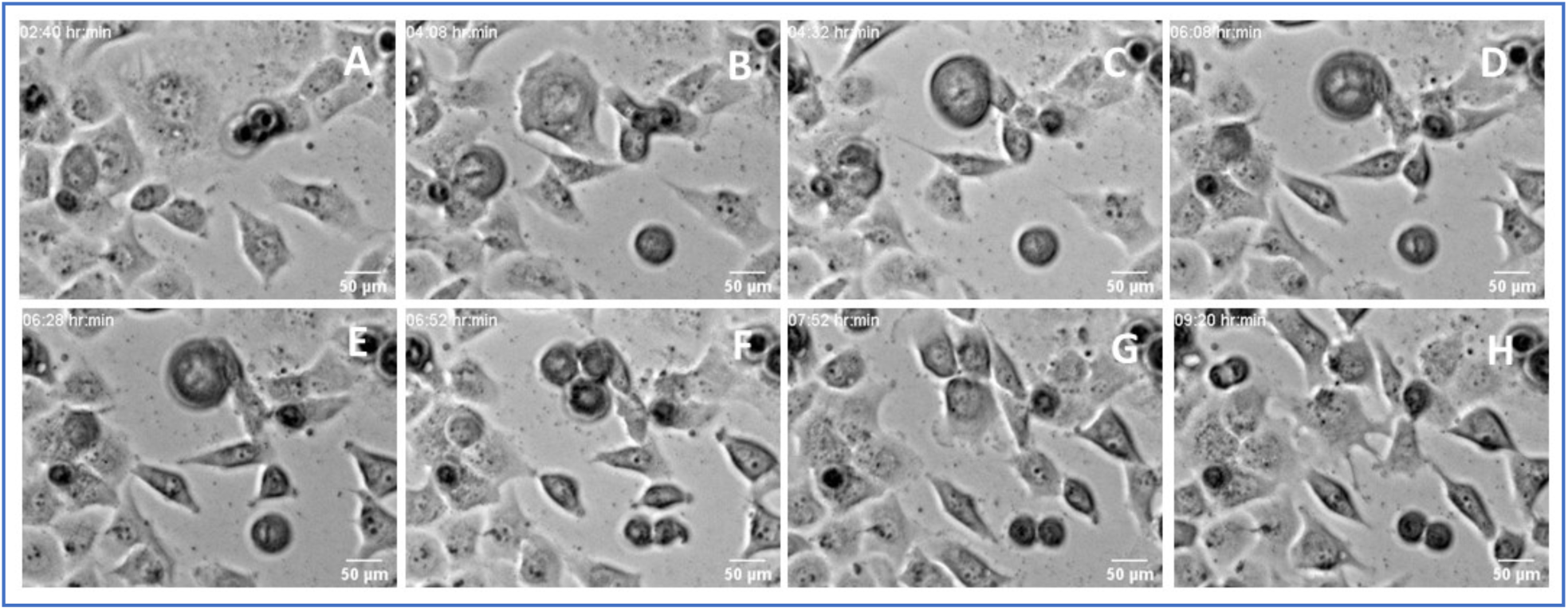
Frames from Supplemental Movie 6: an ALK 4 KO cell undergoes division by three. The large cell in the upper center begins to round up for cell division in panel B. The chromosomes then line up into an unusual configuration, described as a “triskelion,” that is fully formed by panel D. Anaphase is shown in panel E, with the appearance that the chromosomes of each “V” are pulled toward a pole between the arms of each “V”. The resultant daughter cells also spread and began to move away from the centroid of the three, in the directions of the open “V” s seen in panel D.

**Supplemental movies 6 and 7** show examples of cell division into three daughters. **Supplemental movie 7** also shows an interesting pattern attained at apparent metaphase, where the chromosomes line up in a “triskelion.” This pattern was observed in many of the cells that underwent division by three. Sequences from **Supplemental movie 7** are also shown in Figure 8, to emphasize the triskelion, and how that configuration relates to the three daughters produced.Division by >2 daughters, as emphasized in the Introduction, assures a new distribution of the parental chromosomes.

Although cell division by “other-than-two” must shuffle the genome itself, and this phenomenon goes a long way to explaining the many different nuclear configurations seen in multinuclear cells, it does not directly explain the origin of the many very much smaller SAM/PSAR cells. A candidate for the origin of the smallest, simplest cells is a phenomenon we have not, to our knowledge, seen been described before in mammalian cells. It appears to be more of a break up of the parent cell, and a ”hatching out” of small cells from the parent. Examples of this phenomenon are shown in **Supplemental Movies 8 and 9**, and a sequence from **Supplemental Movie 8** is presented in Figure 12. In general, a multinuclear cell that had developed many, very small nuclei such as those seen in Figure 8**, D and N**, ceased motility as though it was about to round up to begin cell division. The cell appeared then to gradually break up into daughters that each surround one nucleus. Oddly, surrounding smaller cells appeared to invade and undermine the mother cell, thus seeming to enhance the breakup process. Finally the newly freed cells rapidly began to undergo cell division themselves, indicating they had already completed S phase before the break up had started. Further work will be required to understand the origin of the SAM/PSAR cells, but this very unusual phenomenon, almost like a “hatching” or a variation on endocytosis, from a mother cell that has reached some sort of critical mass of tiny nuclei, seems an important candidate for that origin.

**Figure 12.**
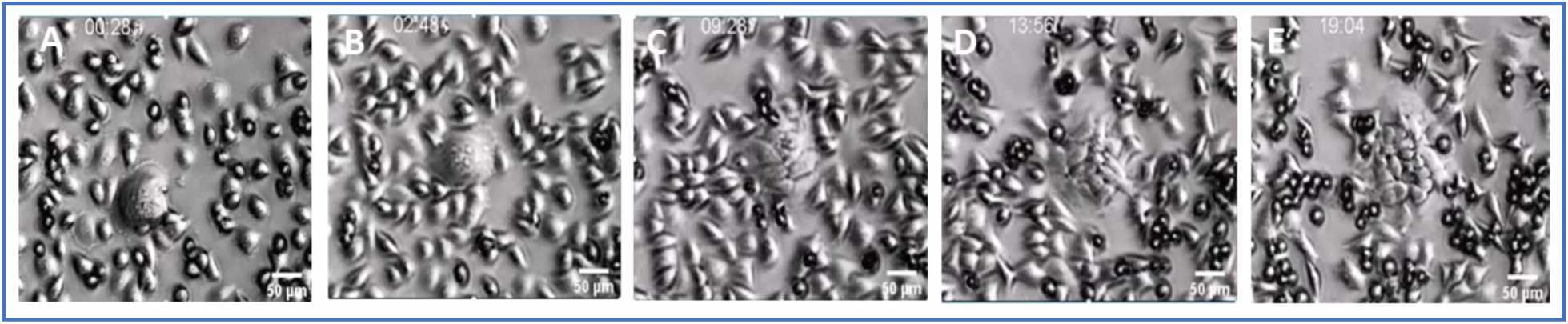
Frames from Supplemental Movie 8: a very large ALK 4 KO multinuclear cell with many small nuclei breaks up into many small cells. The central cells pauses in in movement, and progressively breaks or subdivides into a large number of much smaller cells (panels C-E). The smaller cells then divide by what looks like normal 1:2 cell divisions. This break up has also been described as “hatching”, and seems to have only a weak connection to normal cytokinesis.

**Supplemental Figure 1.**
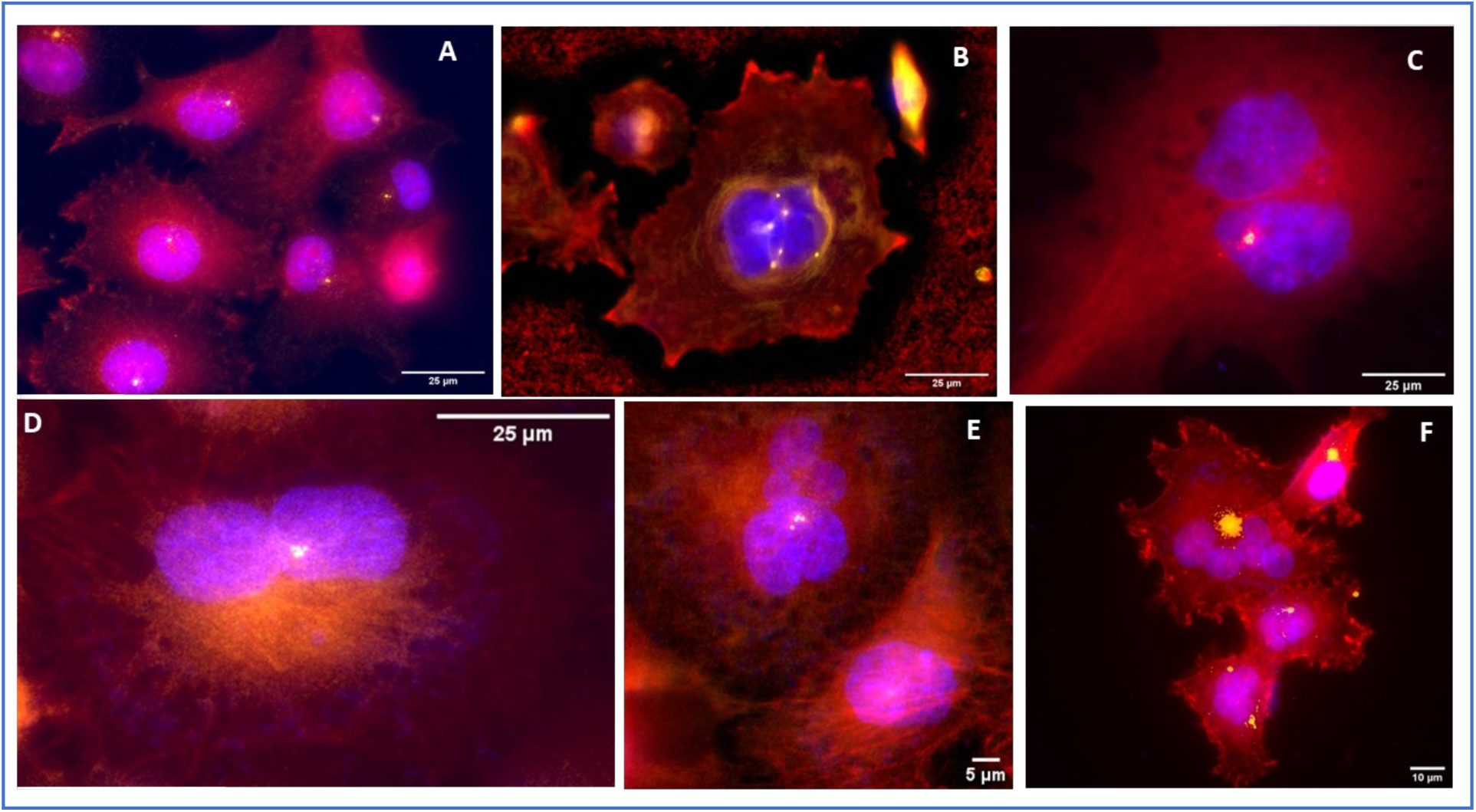
Fixed control and ALK 4 KO multinuclear cells stained for F-actin (red), DNA (nuclei; blue) and for centrosomes (yellow). Antibodies to pericentrin were used to locate the centrosomes in these widefield microscopy images. Control (A) EV cells in interphase displayed the one centrosome/cell expected for this phase of the cell cycle. In contrast, the ALK 4 KO multinuclear cells showed a much more random number and location of these critical organelles (B-F). Note that these images were manipulated in order to enable the visualization of the centrosomes. Nothing was pasted in or moved relative to anything else in the image. The main tool was shade correction and contrast adjustment.

**Supplemental Figure 2.**
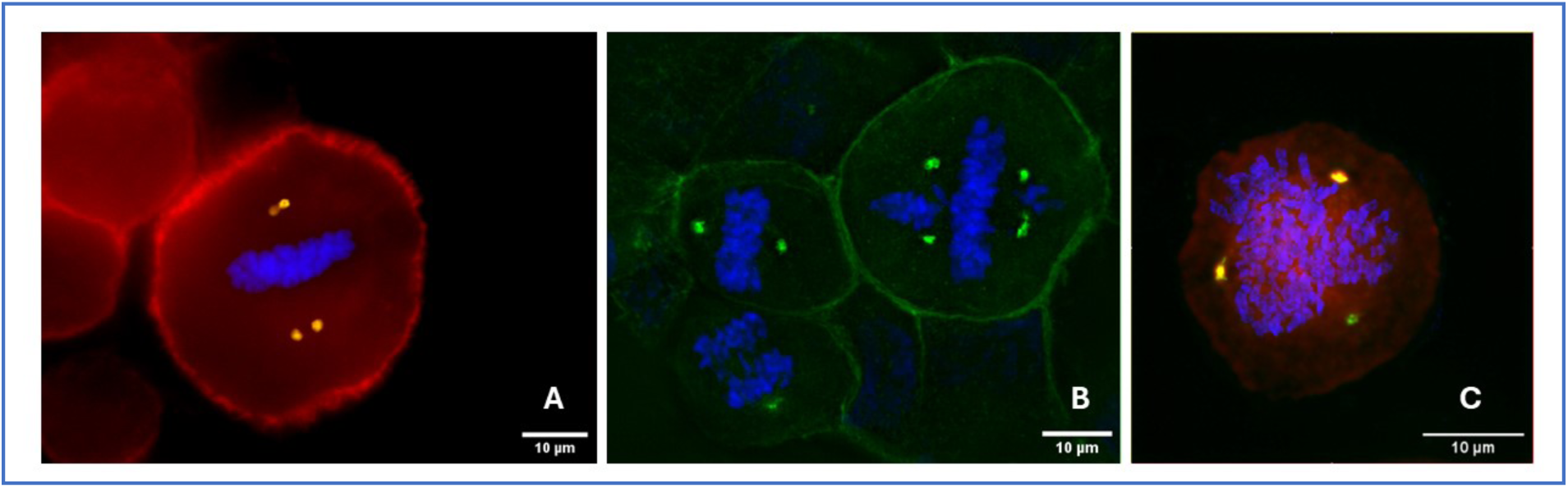
Fixed control and ALK 4 KO multinuclear cells stained for F-actin (red), DNA (nuclei; blue) and for centrosomes (yellow). Structured illumination microscopy was used to image cells that had been blocked in mitosis. Examples of three configurations are shown; each would again shufle the chromosomes in unique ways, but many other patterns are probable.

A factor in the development of the many small nuclei, we well as the many nuclear forms seen in the multinuclear cells, will be the relationship between existing nuclei and the reproduction and position of the centrosomes. As mentioned, Boveri produced unequal cell divisions by introducing extra centrosomes into sea urchin eggs, and intuitively argued that such mismatches might relate to the bizarre nuclear changes he saw in malignant cancers (Boveri 2008). **Supplemental Figures 1 and 2** show fixed cell examples of centrosomes in interphase (**Supplemental Figure 1**) and in metaphase (**Supplemental Figure 2**). Supplemental Figure 1A shows EV cells, which in interphase should have one centrosome, and, in most cases this is the case. **Supplemental Figure 1 C-F** show clusters of centrosomes, or, panel B, centrosomes that are somewhat randomly spaced. **Supplemental Figure 2** shows three examples of metaphase configurations after treatment with nocodazole (MiliporeSigma), fixed and stained as in **Supplemental Figure 2**, but using SIM microscopy to better define the structures at metaphase.

The patterns seen are predicted to further mis-segregate the genome in anaphase. Delineating the patterns of centrosome duplication, movement, and sorting in daughters will be an important factor in understanding and possibly preventing, the genomic degradation that takes place in the multinuclear cells, and thus the many diverse cell types as metastases progress.

## Discussion

Between the time that a mutation stimulates unnecessary cell division in one tissue cell and the much later advent of metastatic disease, lie processes that greatly decrease the survivability of any cancer. A key characteristic of these processes is the development from that one founder cell to descendants with a wide-enough variety of traits that a subset population is sufficiently preadapted to survive the challenge of new tissue environments, immune surveillance, and therapeutic agents. That subset population can then persist and evolve to the detriment of the patient.

We have shown in this one model system that single cells can diversify in previously unsuspected ways to form much smaller, active cells, and also much larger cells with many nuclear configurations. We show that a high proportion of the ALK 4 KO cells have many fewer than 46 chromsomes/cell, and many have very few chromosomes at all. Yet these cells divide normally, grow well in culture, and exhibit properties consistent with being the “seeds” to spread the cancer into new tissues. We show that cytokinesis failure does not necessarily result in senescence of the resulting multinuclear cell, and that cytokinesis failure and merging can explain the origin of multinuclear cells. We show that the multinuclear cells undergo cell divisions of greater-than-two that must result in new arrangements of whole chromosomes into the daughters. Finally, we show an unexpected form of breaking up of a multinuclear cell into many very small cells. This “hatching” appears to be related to an attempt at cytokinesis, as the mother cell appears to begin to round up, but dividing into many cells appears qualitatively different from normal cell division, as is the rapid entry into mitosis of the many daughters. This process seems most likely to be the origin of the smallest cells. Delineating the progression from a simple binuclear cell to large cells with many small nuclei will be critical to understanding how, potentially, this process could be inhibited, thus preventing the generation of the small, metastatic seeds.

Is it reasonable that human cells with so few chromosomes could be viable? Our observations of the smallest cells show that many of the smallest cells are viable, at least for the observation period. Looked at from an evolutionary perspective, viable cells with a much-reduced chromosome number may not be completely surprising. There is good agreement that the chordate lineage underwent two rounds of whole genome duplication early on in our evolution (Meyer and Schartl 1999; Gu et al. 2002; Dehal and Boore 2005). For humans, that would imply a ballpark number of chromosome pairs reduced from 46 to 23 to 11 to 13. Thus, our ancestors made out reasonably well with 12 chromosomes; 6 from each parent. Even more remotely, at the dawn of the last common eukaryotic ancestor (LECA), Goodenough and Heitman (Goodenough and Heitman 2014) argue that our most primitive eukaryotic ancestor would require merging between cells in the ‘experimental stage” of sexual reproduction, cycling between haploid and diploid states. Although the number of chromosomes that LECA had to accommodate is not certain, a number smaller than 6 seems almost certain, and one or two seems the most likely. Thus, it is not surprising that these smallest, most chromosome poor, cells can survive, albeit with a limited behavioral repertoire and limited controls. Moreover, it is intriguing to consider that these cells are an example of genomic degeneration that mimic certain aspects of behavior reflective of very early states of eukaryotic evolution. Such primitive behavior might include the bidirectional tracking behavior seen in Figure 6 and **Supplemental Movie 1**, and the apparent ease with which they merge with other cells. Thus, advanced metastatic cell colonies may include the most degenerate cells, bearing only a faint relationship to the parent tissue or even to mammalian cells. Understanding that this might be a common feature of advanced malignancy, and developing more realistic cell models of metastatic cell colonies, should open the door to finding new vulnerabilities in this complex mix of cells.

If merging behavior is a common but unappreciated behavior of metastatic cells, then the transfer of chromosomes or chromosome fragments to healthy cells must be considered. An obvious example of a chromosome that for pancreas and many GI cancers makes a particular difference is chromosome 12, which contains the gene for KRAS. Cells that received one or more copies of this chromosome, or even just the short arm (12p), would also inherit the drive toward more active proliferation and thus could have a selective advantage. If a SAM-like or PSAR cell that has inherited the part of chromosome 12 with mutated KRAS were to invade or merge with a distant cell in a different tissue, the incorporation of the cancer-promoting gene could, theoretically, destabilize that cell or tissue and cause a secondary carcinoma. In this way the primary cancer would spread not only by metastatic pancreas cells themselves, but by the delivery of a streamlined PDAC genome with cancer promoting genes to the cells of a distant tissue. An unfortunate consequence of this possibility is that the new tumor would be pre-adapted to the new tissue. Depending on how large the PDAC genome was delivered to a new cell, and how much of its genome became incorporated, the resulting metastases would have characteristics of both the invaded and invading cell, creating stimulated chimeric cells within the invaded tissue.

Moreover, markers thought to be specific for a particular carcinoma might then be expressed in the chimeric cells whose origin might then be misidentified. Such cells might also obtain further camouflage from immune surveillance. If this mechanism, or similar, is real, then metastasis could be considered, not only as Hansemann argued to be the wandering and growth of single cells from the primary site, but possibly as the inheritance of an activating chromosome or set of chromosomes. A corollary to this argument is that KRAS inhibitors would be expected to continue to offer benefits to patients even any derived cell subtype of whatever ploidy; even the most degenerate. This suggests that KRAS inhibitors would be effective at constraining the growth cycles of even the most degenerate cells, slowing the progress of even advanced metastatic cancers. Moreover, this effect should be relatively slow to generating resistance, since the cells without mutated KRAS would be much less prone to proliferate. Analogies with other cancers, with oncogenes such as Myc, EGFR, HER2, PI3Kca and so on, may be useful.

On the other hand, approaches that target specific features of PDAC, such as particular antigens for immune therapy, could evolve resistant clones as the cells that lost the chromosome and gene for that antigen would be selected for and proliferate. Chemotherapeutic agents generally block the cell cycle by inhibiting mitotic spindle function, or by inducing DNA damage and stalling DNA synthesis. Unfortunately, with P53 not functioning, the affected cells would often not apoptose or even senesce, but continue to cycle. Moreover, the disruption of normal cycling is likely to produce mitotic or cleavage mistakes, resulting in multinuclear cells or more subtly aneuploid cells. Our present observations strongly suggest that these multinuclear or damaged cells will, once metastatic degeneration has begun, find ways to divide, resulting in further genomic degradation and variety. Thus, chemotherapeutic agents, in the context of non-functioning P53/ apoptotic machinery, can help to cause the genetic diversity that promotes eventual drug resistance. Combining targeted, immune-based, therapies with inhibition stimulating oncogenes, would, by slowing the rate of cell cycling, also slow the rate of cell degradation and thus potentially reduce the rate of production of resistant clones.

In contrast to the argument that the inhibition of stimulating oncogenic mutations would be beneficial at all stages of cancer, our model suggests that a restoration of p53/p16/Rb function, while it might be beneficial in very early stages of metastatic development, would not be useful once the smaller cells have been generated. These cells were consistently able to divide one-to-two, and would therefore not be subject to any normal checkpoints.

In mammals, cells with extra, sometimes many extra, chromosomes are expected to have exited the cell cycle and not divide. But there are exceptions (Davoli and De Lange 2011). Megakaryocytes, placental parietal trophoblast giant cells, and skeletal muscle cells, all become polyploid during normal development, and even cardiac myocytes have been found to be over 50% multinuclear (Buttitta and Edgar 2007; Deng et al. 2017; Windner et al. 2019; Bergmann et al. 2015; Lan et al. 2025). An especially intriguing case has recently been demonstrated, that liver hepatocytes are very often bi-nuclear. Hepatocyctes are able to undergo successful cell division with two nuclei present, and do not degrade their genomes (Knouse 2025). Moreover, this appears to be a benefit to rapid repair of a damaged liver. In addition, many plant and even other vertebrates typically are tetraploid, and, again, are able to grow and reproduce successfully. Understanding the strategies by which polyploid cells are generated in the first place, whether through fusion or endoreplication, and the mechanisms that successfully control mitosis and cell division to maintain genomic integrity, may offer clues to how to minimize the shuffling and degradation of the human genome by continued cycling of very polyploid cells in cancer.

Many cancer researchers have pointed to the contribution of the tumor microenvironment (TME) to cancer progression (Lafaro and Melstrom 2019; Henke et al. 2020; Anderson and Simon 2020). This is pictured to be comprised of cells from the tissue that is being invaded, plus fibroblasts, immune cells and deposition of complex extracellular matrix. The present demonstration of diversification from single cell clones, and the appearance of cooperation of subtypes that literally support the smallest, weakest, least adherent cells, suggests that a metastatic cell can essentially create its own cellular microenvironment. Such self-generated colonies could already contain an equivalent of cancer-associated fibroblast (CAF) cells in SAM-like cells, and these fibroblast-like cells would inherently be predisposed to carry smaller ‘sibling’ cells along with them into new tissue. The potential that diverse descendants of a primary tumor cell could by themselves form a cooperating, diverse, tumor microenvironment adds complexity to the interpretation of TME, as did the phenomena of cell merging discussed above, and suggests that finding ways to interfere with this type of cooperation might, in itself, be generally useful in slowing down metastatic disease.

It is possible that these observations are unique to pancreas cancer cells, or to Panc 1 cells in particular after ALK4 loss. We mentioned above that the Panc 1 cell line appeared poised to replicate the statistical pattern of destabilization seen in most PDAC, with KRAS mutated, P16 missing, and SMAD 4 present. The crispR KO of ALK4, which signals downstream through SMAD4, appears to represent the loss of a final mechanism that had prevented the continued cycling of bizarre multinuclear cells, thus allowing the generation of the many surprising phenomena described herein. So, in the sense that this specific pattern of gene activation and loss, coupled with the loss of ALK4, is unique, the present observations may offer a serendipitous, clear-cut example of processes that are generally difficult to capture.

On the other hand, the fact that a similar pattern of stimulating mutation, slow development with progressive nuclear disorganization, and then rapid appearance of adaptable, less curable metastases is common for most solid tumors, argues that our observations may be somewhat common in other carcinomas and cell lines derived therefrom. Since not all tumor-derived cell lines will have the same pattern of gene alterations, not all cell lines would be expected to exhibit the same patterns. Rather, like Panc 1 cells before ALK4 KO, they may be in various stages along the same general path. By optimizing cell culture procedures to preserve smaller, poorly attached cells, observing the buildup or shedding of cells, taking note of heterogeneity in the culture, and observing a high percentage of multinuclear cells, other cell lines could be assessed for where they are on the same trajectory. Preliminary observations in our laboratory showed some aspects of this phenomenon in other cell lines with or without KO of ALK4, in terms of higher frequencies of multinuclear cells, and rarely the evolution of smaller, less adherent cells. It is hoped that corroborating observations in other cells lines will allow these phenomena to be studied in the depth they deserve.

## Methods

### Cell Culture

Panc1 human pancreatic cell lines were obtained from ATCC and each was cultured in DMEM medium, supplemented with 10% FBS and 1% penicillin/streptomycin. The ALK4 WT (Cat# 80879) and ALK4 CA (Cat# 27223) DNA plasmids were purchased from Addgene.

The CRISPR ALK4 knockout cells were generated using pLentiCRISPRv2-puro vector (Addgene, Cat# 98290). Three gRNA sequences (Supplemental Table) were annealed separately to BsmBI-digested pLentiCRISPRv2-puro vector using T4 ligase. The insertion of gRNA was confirmed using Sanger sequencing. 293T cells were transfected with the final pLentiCRISPRv2-puro-gRNA vector along with the psPAX2 (Addgene, Cat# 12260), and pMD2.G plasmids (Addgene, Cat# 12259), and the resulting lentiviral condition medium supplemented with polybrene (Sigma, Cat# TR-1003-G) was cocultured with a parental Panc1 or MDA-MB-231 cell lines followed by puromycin selection (1 μg/ml). The ALK4 KO clones were validated using Sanger sequencing and western blotting.

Cells were grown to 60-80% confluence before passaging using 0.05% trypsin/EDTA (Gibco) and replated at 20-30% confluence. Brief, gentle, washing in PBS before trypsinization helped preserve less adherent cells when present, but some are unavoidably lost. Gently swirling the culture plate in those that had not overgrown and collecting the media before washing, centrifugation of the media, allowed the collection of the less adherent cells when needed.

### Brightfield Microscopy

Routine cell culture microscopy was carried out using a Zeiss Axiovert microscope equipped with a phase optics and an ocular ToupView digital camera. Images acquired at 4X at 2592 x 1944 pixels, 1.2 px/µm.

### Nuclear volume quantitation with widefield fluorescence microscopy

EV control and ALK 4 crispR KO Panc 1 cells were plates on #1.5 22x22 coverglass squares in 6 well plates and grown until 60-80% confluence in normal 10% DMEM medium. Medium was then replaced with PBS, then with 4% formaldehyde for 5 min, gently washed, and then stained with 1 µg/ml DAPI (4ʹ,6- diamidino-2-phenylindole; Fisher Scientific). 61 images from three experiments images were then acquired using a Nikon 60 xPlan Apo 1.40 Oil Ph3 objective and Nikon TE 2000 inverted microscope, captured with a Photometrics CoolSnap CCD camera (1392x1040 pixels), using MicroManager software (doi:10.14440/jbm.2014.36). Tabulation of estimated nuclear volumes for the ≈200 cells for each condition was carried out using FIJI (doi:10.1038/nmeth.2019). The oval or surround tool was used to evaluate area, length and width. Volumes were estimated as area at best recorded focal plane times ½ the oval width as a surrogate for the radius. The histogram was generated starting at 50 cubic microns, and binning at increments of 150 cubic microns.

### Flow Cytometry

Panc 1 KO and EV cultures were plated and grown to similar confluences on tissue culture plastic, trypsinized, pelleted and fixed in parallel with ice cold 70% ethanol. Cell pellets were mixed into a propidium iodide (50ug/ml) solution in FACS buffer (0.1% BSA in PBS with 0.1% RNAase) and kept in the dark at room temperature for 30 min. The cells were then pipetted past 35 µm strainer caps into flow tubes and kept on ice in the dark until analyzed on the BD Canto cytometer using FACSDiVa software. The cell cycle was analyzed using FloJo and ModFit software.

### Mitotic Chromosome Squashes

ALK 4 Ko cells were grown in normal 10% DMEM in 10 cm tissue culture plastic until ≈50% confluent, and then the medium gently exchanged for Medium with 50 ng/mL nocodazole (MiliporeSigma). Cells were grown for 10-12 hrs in nocodazole, and then prepared for cell squashes. Importantly, cells that round up to begin mitosis are poorly adherent, so we gently swirled the dish and collected the supernatant. The cells were then spun down (clinical centrifuge, 400xg, 5 min) and the pellet added to the main cells collected after trypsinization. The remaining adherent cells were placed in 0.05% trypsin/EDTA (Gibco; Thermo Fisher Scientific) for 5 min, collected in 10 mL centrifuge tubes, spun down as before, and the pellets collected. Pellets were gently resuspended in PBS, added together, and gently centrifuged in PBS to remove supernatants. The pellets were then gently resuspended in hypotonic medium (0.56% potassium chloride in distilled water) at 37 deg for 30 min with gentle rocking motion, centrifuged to remove swelling buffer, and the pellet gradually resuspended in fixative (3:1 methanol:acetic acid), then agitated with pipetting to keep cells suspended. The cell suspension was placed on ice for 30 min, centrifuged again, resuspended in fixative for 30 min, and repeated once more until a slightly opalescent suspension was reached. Microscope slides (1x3 in) were soaked in ice cold fixative, dried with kimwipes, and then small volumes of the cell suspension were dropped onto the slides, and the drop rapidly dried on the slide. Repeated drops were added, the slide dried, and the slide preserved with Vectashield with DAPI (Vector Labs). Slides were viewed in widefield fluorescence for DAPI.

### Time lapse microscopy

Motility and behavior of the EV and Ko cultures was carried out in 12 well tissue culture plates (Corning, Inc) plates in a Zeiss Live Cell Station with CO2 and temperature control. Images were taken once per 4 minute intervals, which allowed a survey of many areas of interest during the same time interval. Imaging was generally with a 5x objective in phase contrast mode. This was sufficient to observe large areas at one time, and with a resolution (1392 x 1040; Photometrics CoolSnap) sufficient to show the behavior of individual events of interest. Cultures were typically observed for several days at a time.

### Immunofluorescence microscopy

Immunofluorescence microscopy was carried out as described for widefield fluorescence microscopy, above, but with a brief treatment with 0.2 % triton X100 to permeabilize the cells before addition of 1% antibodies to pericentrin (ABclonal Inc, A11629, Rabbit polyclonal). For widefield imaging of centrosomes, fields of view showing multinuclear cells were searched for, while for SIM imaging, cells in mitosis were selected.

**Structured illumination** images were collected at the Duke Light Microscopy Core on a Zeiss ELYRA 7 with laser excitation at 405 nm, 488 nm, and 561 nm (all at 1%) and emission split to LP560 (camera 1) and SP560 (camera 2) in two sequential tracks using dual pco.edge 4.2 sCMOS cameras. Lattice SIM images were collected with a pattern grating size 32um and 13 phases and a camera exposure time of 50ms. Raw datasets were processed using SIM Processing in Zen Software with “adjusted” setting “normal to baseline cut”. Z-stack images were collected using a 63x plan apochromatic 1.4 NA oil objective attached to a Zeiss Observer 7 SR inverted stand with an X,Y linearly encoded Marhauzer motorized stage and piezo Z stage with 500µm range set to 110 nm z-step size.

## Acknowledgments

Thank you to Lisa Cameron, Light and Microscopy director at Duke.

Thank you to Manqi Zhang Department of Medicine, Division of Medical Oncology, Duke University Medical Center. She contributed to sample preparation, culture maintenance, experimental execution, and data interpretation.

Thank you to Gerard C. Blobe, also of the Department of Medicine, Division of Medical Oncology, Duke University Medical Center. He supervised the project, provided laboratory resources, and secured funding and institutional support.

This manuscript represents the final scientific work of E. Tim O’Brien III. This paper’s collaborators gratefully acknowledge his perseverance in completing this study during his fight with pancreatic cancer. We dedicate this publishing effort to his contributions to the field and those he inspired.

## Author contributions

ETO conceived and designed the study, performed experiments, acquired data, conducted initial analyses, analyzed and interpreted the results, and wrote the manuscript.

## Supplemental Movie Legends

Movies published to figshare, DOI 10.6084/m9.figshare.30782168

**Legend to Supplemental Movie 1.** Colony formation by ALK 4 KO Panc 1 single cell clones showing diversity in morphology and behavior. Cells were plated through 8 µm pore uncoated transwell filters (Corning, inc) directly onto 12 well multiwell tissue culture plastic that had been treated with 0.01 mg/mL Matrigel (no growth factors). Normal 10% serum medium was present on both sides of the filter. After 24-48 hrs the filter wsa removed, medium replaced, and the cells allowed to grow for 14 days. Time lapse recordings were obtained using a Zeiss Live Cell Station with 5% CO2 and 37 deg at intervals of 4 min. Images were at 5X brightfield to a CoolSnap 1392x1040 pixel CCD camera (Photometrics; Roper Scientific).

**Legend to Supplemental Movie 2.** ALK 4 KO cells that were able to pass through 3 µm transwell-type filters (Genesse Scientific) directly onto Matrigel- coated tissue culture plastic, were imaged at 4 min intervals, as described for Supplemental Movie 1. They showed motility and poor adhesion once cell division had take place.

**Legend to Supplemental Movie 3.** Large motile multinuclear cell shows rapid motility, and the property of transporting much smaller cells for long distances. Cells were grown from cells that had passed through the 3 µm filters, and imaging as described in Supplemental Movie 1.

**Legend to Supplemental Movie 4.** Two cells (yellow arrow) appear to merge together and proceed as one cell. ALK 4 Ko cells that had passed through 3 µm filters were imaged as described in Supplemental Movie 1.

**Legend to Supplemental Movie 5.** ALK 4 Ko cell rounds up to begin mitosis, pauses for several hours, and then does not finish cytokinesis and moves away as a binuclear cell. Imaging was carried out described as for Supplemental Movie 1. Slices from this movie are presented in Figure 10.

**Legend to Supplemental Movie 6.** ALK 4 Ko cell in center of frame round up and divides into three daughters. The daughters then move away independently. Imaging was carried out described as for Supplemental Movie 1.

**Legend to Supplemental Movie 7.** Movie basis for Figure 11; Large cell divides into three daughters. Interesting also for the “triskelion” configuration of chromosomes achieved at one point in the process. Imaging was carried out described as for Supplemental Movie 1.

**Legend to Supplemental Movie 8.** Movie basis for Figure 12; Large ALK 4 KO multinuclear cell with many small nuclei, breaks up into many smaller cells in a very aberrant version of cytokinesis. Many of the daughters then begin to undergo cell division. Imaging was carried out described as for Supplemental Movie 1.

**Legend to Supplemental Movie 9.** A second example of an large multinuclear cells breaking up into many smaller cells without undergoing a standard cytokinesis process. This movie was obtained at 10X phase contrast initial magnification, but otherwise the same time lapse conditions as Supplemental Movie 1.

